# Still Lost in Definitions: How ‘Resilience’ Is Used in Ecology

**DOI:** 10.64898/2026.06.03.729936

**Authors:** Hung-wei Lin, Sruthi M. Krishna Moorthy, Andrew Hector, Roberto Salguero-Gómez

## Abstract

Resilience is a central concept in ecology and environmental policy, yet its meaning and quantification remain inconsistent across subfields. Clarifying how resilience is defined and measured across the subfields of ecology is therefore a critical step towards delivering coordinated efforts to strengthen resilience research. Here, we analyse 594 studies published between 1977 and 2025 to determine how resilience is quantified across ecological contexts. Using large language models to extract structured data and conditional inference forests to assess predictors of metric choice, we show that resilience is most commonly (∼25%) quantified using recovery rate and recovery degree, but no single metric dominates. Crucially, study attributes like organisational level, methodological approach, and disturbance regime explain only a small fraction of variation in metric selection. Despite this apparent inconsistency, more than 90% of studies draw from a shared set of six quantitative dimensions of resilience. This combination of weak constraint and latent convergence suggests that resilience metrics function as a flexible but implicitly standardised toolkit rather than as context-specific constructs. We argue that this hidden structure provides a foundation for a unified, multidimensional resilience framework that can support synthesis across ecological systems and improve the translation of resilience science into conservation and policy.

## Introduction

Global change is accelerating biodiversity loss^1,2^ and altering ecosystem functioning worldwide^3,4^. In this context, resilience has become a central concept in ecology^5,6^ and a cornerstone of environmental policy^7,8^. Yet, despite its widespread use, resilience remains difficult to define and even harder to measure consistently across studies^9,10^. This lack of clarity limits our ability to compare results^11^, synthesise evidence^12^, and translate ecological understanding into actionable management strategies^13,14^.

Since its introduction by Holling^15^, resilience has been interpreted in multiple ways. Early formulations emphasised the capacity of a system to absorb disturbances without shifting to an alternative stable state^16^, highlighting nonlinear dynamics and regime shifts^17^. Subsequent work introduced “engineering resilience”, the rate at which systems return to equilibrium following disturbance^18^, reflecting a more tractable and operational perspective in conservation and management contexts^19^. Over time, additional interpretations have proliferated, spanning multiple levels of biological organisation and focusing on distinct aspects of system dynamics^20^. As a result, resilience is now quantified using a wide range of metrics, from recovery rates^21^ and degrees of return^22^, to measures of resistance^23^, invariability^24^, and tipping-point proximity^25^.

Today, resilience is widely invoked throughout ecology and operationalised in divergent ways^10,26^. Ecologists working across levels of biological organisation, from individuals^27^, populations^21^ to communities^23,28^, ecosystems^25,29^ and landscapes^30^, employ diverse metrics to capture different aspects of system responses to disturbance. For instance, community ecologists may assess interaction-network resilience as robustness to secondary extinctions^23^, or quantify compositional resilience as the persistence or recovery of species assemblages^28^, whereas ecosystem ecologists may evaluate functional resilience, either as the maintenance of photosynthetic performance under stress^29^ or as the temporal stability of primary productivity dynamics^25^. The diversity of foci used by ecologists highlight how resilience is a multidimensional and context-dependent concept. However, while the need to understand how disturbances shape natural systems across levels of biological organisation is increasingly recognised^5,31^, the conceptual plurality of resilience has sown widespread and persistent confusion, hindering integration of resilience research across subfields^9,32^. Previous attempts to unify the concept have largely focused on theoretical clarifications^9,14^ or on advocating specific metrics^10,33^. However, these efforts have not resolved the diversity of practices observed in empirical research. A key unresolved question is whether this diversity reflects meaningful differences among ecological systems and research contexts, or whether resilience metrics are selected more flexibly, independent of study design.

Here, we address this question through a large-scale synthesis of resilience research in ecology. We analyse 594 studies published in leading journals between 1977 and 2025, using large language models to extract standardised information on how resilience is quantified and the contexts in which it is studied. We then test whether study attributes (*e.g.*, level of biological organisation, methodological approach, disturbance characteristics, and measured response variable) predict the choice of resilience metric. Our results reveal a striking pattern: although resilience is quantified in diverse ways, metric choice is only weakly structured by study context. At the same time, the vast majority of studies draw from a common set of metrics to quantify resilience, indicating an underlying convergence in practice. We interpret this combination of weak constraints and shared structure as evidence that resilience metrics function as a flexible but implicitly standardised toolkit. Building on this insight, we argue that a unified, multidimensional framework of resilience is both feasible and necessary to advance synthesis across ecological systems and to strengthen the application of resilience science in conservation and policy.

### Patterns and temporal trends in resilience metric use

Resilience metrics cluster into six shared ways of quantification despite terminological diversity. To enable consistent comparisons across resilience studies, we first standardised the diverse terminology used to quantify resilience in the literature. Reported metrics were harmonised into six quantitative categories—recovery rate, recovery degree, recovery time, resistance, latitude and invariability—each assigned to a single term (Table 1). Importantly, we note that a given study may employ multiple metrics. We find that resilience research is dominated by direct-response approaches, yet exhibits substantial heterogeneity in how resilience is quantified. Indeed, across the 594 papers examined from 15 journals between 1977 and 2025, 80.8% quantify resilience based on directly observed system responses to disturbances, whereas 19.2% studies rely on inference-based approaches using proxy indicators such as interaction strength or land-use types (Fig. 1). Of the studies examining direct-responses to disturbances (n = 479), we identify at least 21 distinct combinations of resilience metrics. Here, recovery rate and recovery degree are the most frequently used categories, occurring in 32.8% and 31.3% of studies, respectively. Recovery time (17.5%) and resistance (14.4%) are also common, whereas multidimensional approaches combining multiple dimensions of resilience remain rare (9.2%).

**Table 1.**
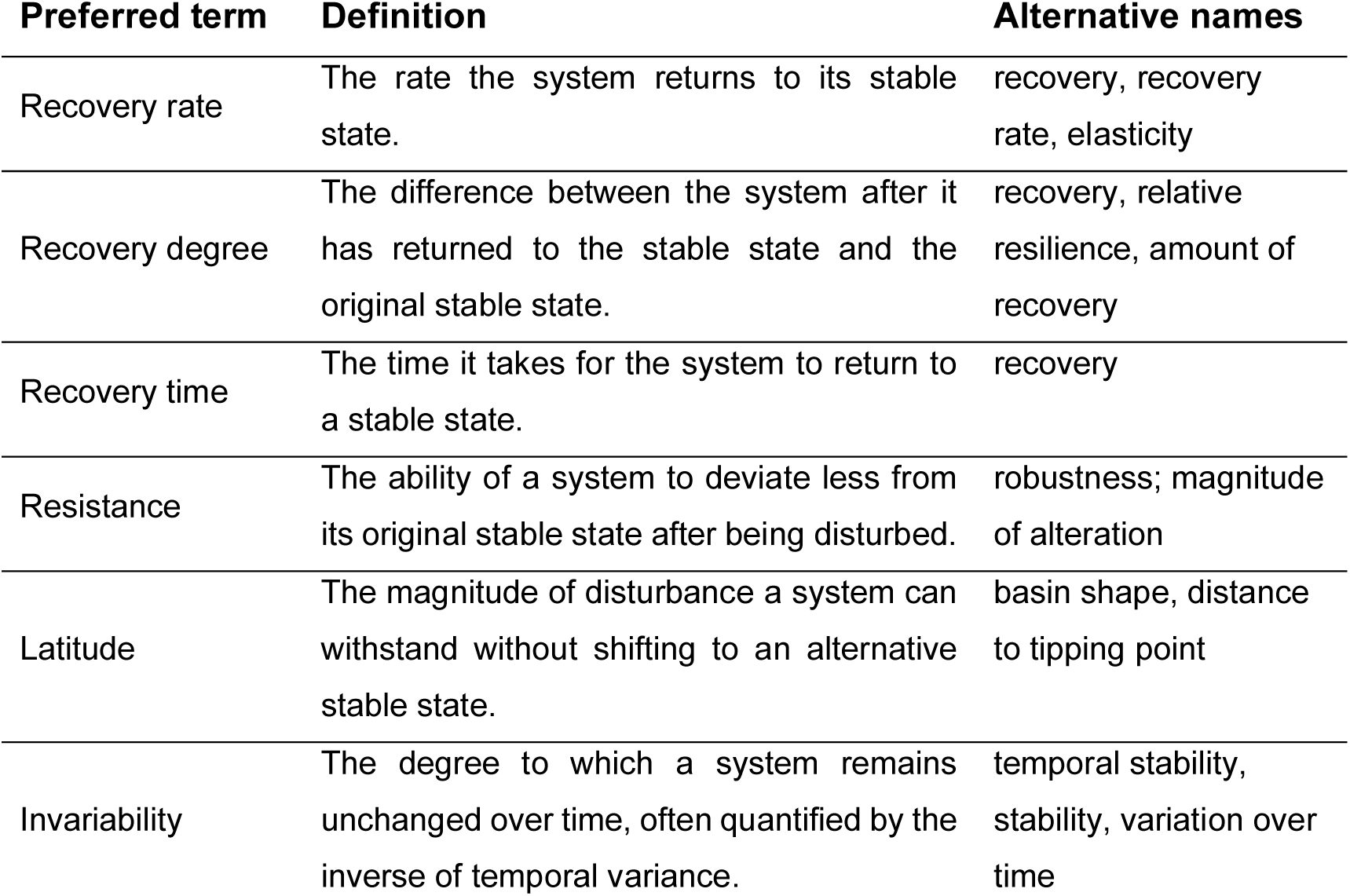
Definitions and standardised terminology for the six common components of resilience quantification. The table harmonises diverse terms in the literature by assigning one preferred name to each component and listing a concise definition, and frequent alternative names found in our dataset. Metrics that did not align with these categories were classified as “other”. Recovery degree (extent of return), recovery rate (rate of return), and recovery time (time to return) are treated as distinct dimensions. In this study, a single sample (*i.e.*, a study quantifying resilience) may have multiple metrics to quantify resilience. Representative examples of studies corresponding to each category are provided in the Supplementary Methods.

**Figure 1.**
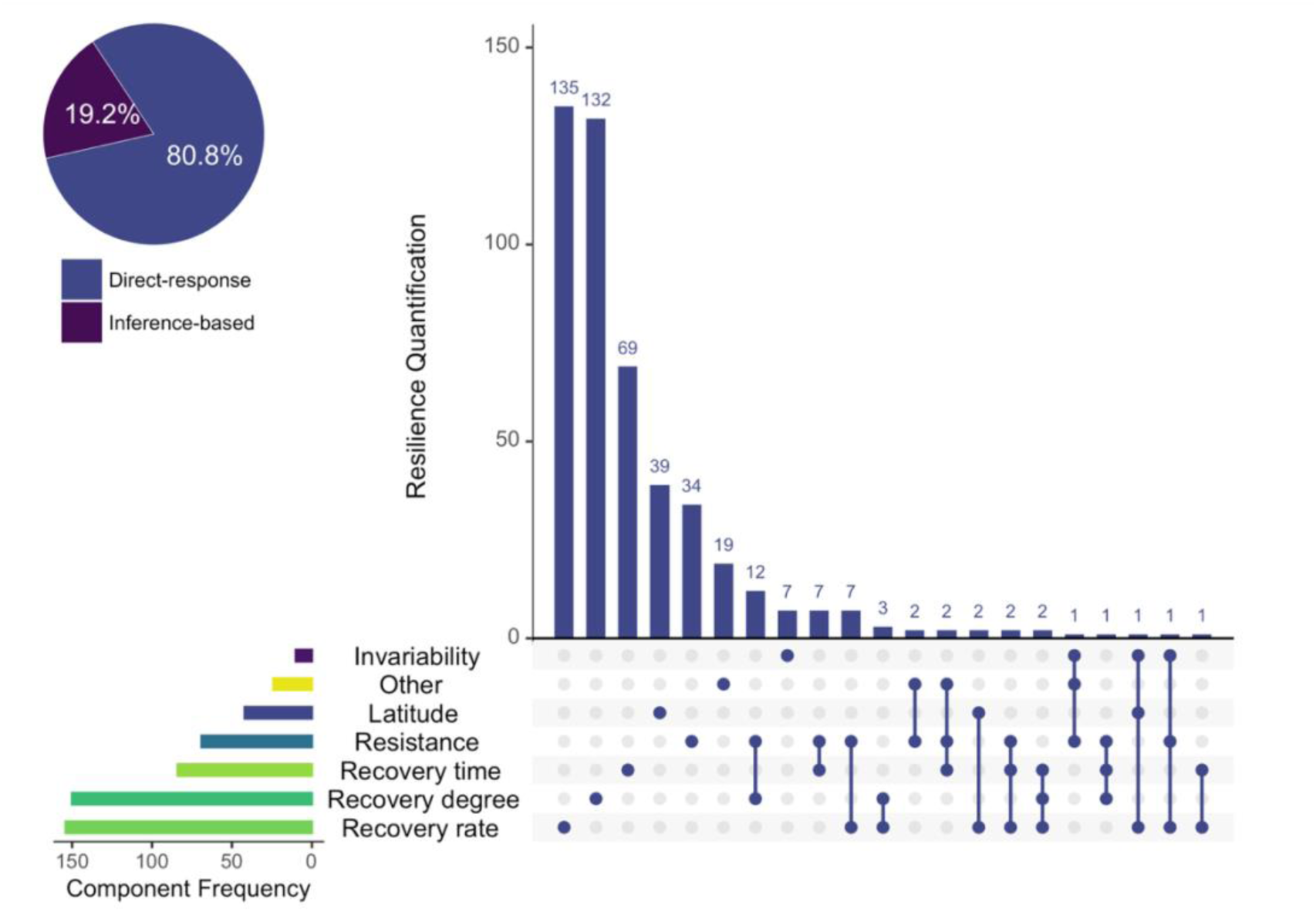
Frequencies and combinations of resilience metrics in ecological studies across 15 high-impact journals. The figure summarises the frequencies and co-occurrences of resilience metrics across 594 peer-reviewed publications from 15 selected journals (Supplementary Table 7). The pie chart (upper left) shows the proportion of studies by measurement basis: direct-response studies, which quantify resilience from observed system responses to disturbance, and inference-based studies, which infer resilience from proxy indicators or models rather than from direct system responses. The UpSet plot (right) shows how the six most frequent quantitative categories (recovery rate, recovery degree, recovery time, resistance, latitude and invariability; definitions in Table 1) co-occur in direct-response studies; all remaining metric types are grouped as “other”. Vertical bars represent the number of studies for each metric combination. Filled dots connected by lines indicate which metrics are included in each combination. Horizontal bars on the left show the overall frequency of each metric category; bar colours correspond to the legend.

Despite a rapid increase in resilience research, we find no evidence that the field is converging on a common metric (Fig. 2). Even in the most recent decade, no metric exceeds one quarter of usage. To examine whether the distribution of resilience metrics shifts with the development of the field of ecology, we analysed temporal trends across five-year intervals. Across the entire study period, recovery rate and recovery degree consistently remain among the most frequently adopted resilience metrics, despite the total number of studies explicitly quantifying resilience increases sharply over time. However, even when combined, these two metrics account for less than half of all quantifications during the most recent decade (2016–2025). Two temporal shifts are particularly evident. First, inference-based approaches began to increase during 2011–2015 and now account for approximately one quarter of resilience studies as per the most recent period (2021–2025). Second, the multidimensional quantification of resilience emerges during 2016–2020 and subsequently stabilises at around 10% of studies. Aside from these changes, the proportional representation of other resilience metrics remains stable over time. Focusing on the most recent period (2021–2025), no single metric category exceeds one quarter of usage.

**Figure 2.**
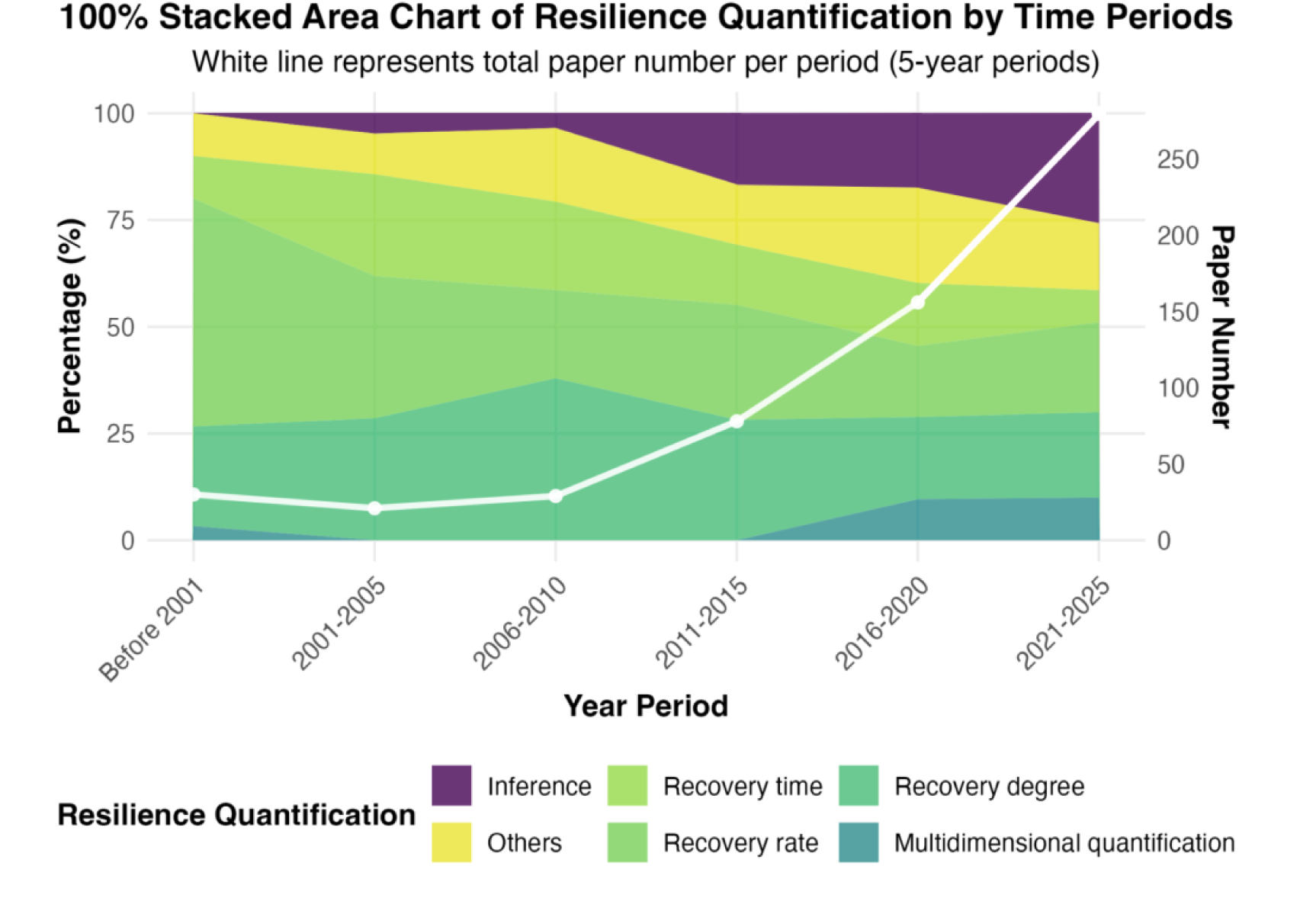
Changes in the relative frequencies of resilience-quantification categories over time (1977-2025). The 100% stacked area chart shows the relative frequencies of resilience-quantification categories in peer-reviewed publications by five-year periods (pre-2001 pooled). Publication years are binned into six periods: pre-2001, 2001–2005, 2006–2010, 2011–2015, 2016–2020, and 2021–2025 (x-axis). Colours indicate categories: inference, recovery time, recovery rate, recovery degree, multidimensional quantification, and others. “Inference” denotes studies that quantify resilience from proxy indicators or models rather than from observed system responses to disturbance. Definitions of the three recovery categories appear in Table 1.“Multidimensional quantification” refers to studies that quantify resilience using multiple metrics jointly (*e.g.,* combining recovery degree with resistance). “Others” pools less-frequent quantification types not listed above (*e.g.,* resistance, latitude). Area heights correspond to within-period percentages (left-hand y-axis), which sum to 100% per period. The white line plots total paper counts (right-hand y-axis).

### Distribution of study attributes

Our results show that resilience research spans multiple levels of biological organisation, methodological approaches, and measured variable types, with numerous combinations represented across studies (Fig. 3). The categories of measured resilience variable types are defined in Supplementary Table 1. Most studies focus on ecosystem (36.5%, n = 217) and community levels (35.0%, n = 208), whereas investigations of resilience in populations (14.1%, n = 84), landscapes (9.1%, n = 54), and individuals (5.7%, n = 34) are less frequent. Field observational studies predominate the landscape of resilience research (58.2%, n = 346), followed by model-based simulations (30.1%, n = 179) and field experiments (25.8%, n = 153). Significant associations emerge among the levels of biological organisation, chosen methodological approach, and measured variable type (Supplementary Table 2). For instance, landscape-level studies are more likely to employ model-based simulations and to quantify resilience by measuring environmental context variables, such as land-use type or network properties. Ecosystem-level research predominantly measures functional indicators, including productivity and biogeochemical fluxes, whereas community-level studies emphasise structural attributes such as composition and diversity and more frequently rely on field experiments. Population-level studies most often quantify abundance-based measures and process-based parameters such as population growth rate and recruitment rate, while these studies show a greater representation of model-based simulations. In contrast, individual-level resilience investigations primarily focus on physiological or growth-related indicators.

**Figure 3.**
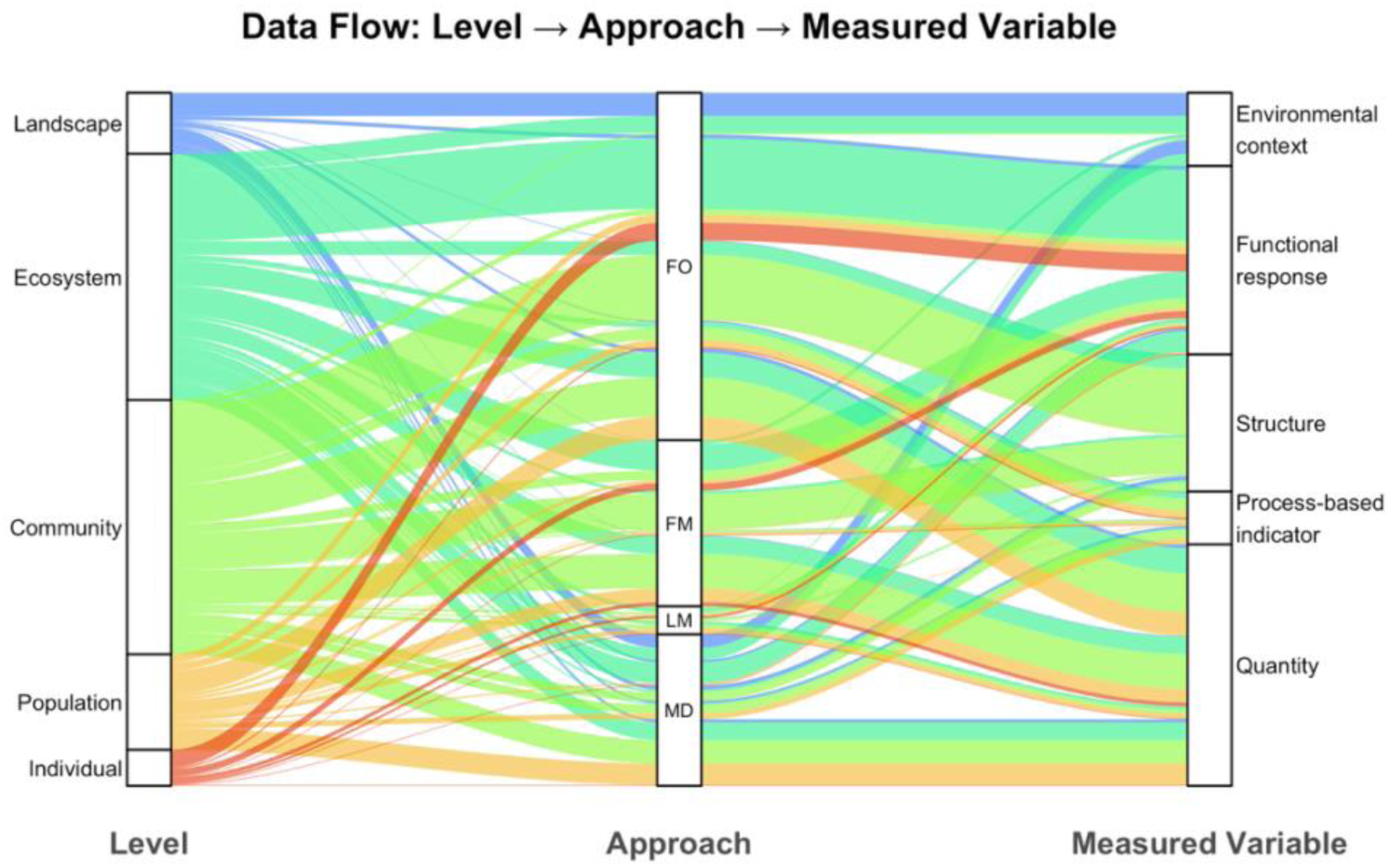
Associations among biological level, methodological approach and measured variable type in resilience-quantified studies. The Sankey diagram shows connections between levels of ecological organisation (left), methodological approaches (middle), and measured variable types (right). Levels include individual, population, community, ecosystem and landscape. Approaches comprise field observation (FO), field manipulation (FM), laboratory manipulation (LM), and model-based simulation (MD). Variable types include environmental context, functional response, structure, process-based indicators and quantity (definitions in Supplementary Table 1). Link width is proportional to the number of study-level occurrences. Colours correspond to the level of origin (left axis). Because individual studies may have multiple levels, approaches or measured variable types, entries were expanded such that each distinct adjacent pairing (level–approach and approach–variable) contributes one link. Totals therefore exceed the number of studies.

The context of the disturbance under examination, such as its pattern, type, and duration, also exhibits a non-random landscape (Supplementary Fig. 1; Supplementary Tables 3 and 4). The categories of disturbance type are summarised in Supplementary Table 5. Among the most striking patterns here are the greater representation of pulse (short-term, discrete events^34^) relative to press disturbances (long-term, continuous pressures^34^) in studies that manipulated disturbances, and the lower occurrence in unmanipulated studies of disturbances involving changes in resource availability or chemical conditions, such as nutrient limitation or pollution. The duration of the study also varies significantly across disturbance types, with studies examining climatic, geophysical, and fire disturbances generally being characterised by longer observation periods than those focused on structural, chemical and resource availability disturbances (Supplementary Table 4).

Among the studies based on field data, resilience research is concentrated in North America, Western Europe, and China (Supplementary Fig. 2). Of the recorded disturbance events, climatic disturbances constitute the largest share (39.5%), reflecting the frequent study of phenomena such as drought and temperature extremes. Other disturbance categories are less frequently examined, such as biotic disturbances (13.5%) or fires (8.8%). Furthermore, using the IUCN habitat classification^35^, we show how forest ecosystems account for the largest share (23.5%) of studies examining resilience globally, followed by neritic marine systems (16.6%), grasslands (13.1%), and wetlands (12.8%). In contrast, the study of resilience in deserts, savannas, and urban systems contributes together only 6.9% of the overall share (Supplementary Fig. 2c). Moreover, taxonomic coverage in the examined publications is strongly skewed toward plants (51.9% of studies that specify focal taxa), with animal resilience being examined in 23.7% of such studies and other kingdoms appearing less than 6% (Supplementary Fig. 3).

### Study attributes as predictors of resilience metric choice

Study attributes exert limited but detectable influence on the choice of resilience metric. To assess this relationship, we fit conditional inference forests to predict the resilience metric category from eight study attributes (see Methods). Classification performance of our model exceeds the expectation by chance (mean macro–F1 = 0.293, 95% CI: 0.221–0.363; compared with a random baseline of 0.167 for six classes), indicating that study context provides only limited explanatory power for the choice of resilience metric, although overall discriminative power remains modest. The prediction accuracy of our model varies across resilience metric categories (Supplementary Table 5), implying that certain metric categories exhibit particularly weak associations with study attributes. For instance, the quantification of resilience via multiple metrics is the most difficult category to classify correctly (F1 = 0.029), followed by recovery time (F1 = 0.196), which is often confused with recovery degree and recovery rate (Supplementary Fig. 4). Together, these results indicate that resilience metrics are not strongly constrained by ecological context.

Our permutation importance analysis identified the response variable measured in the resilience study as the most influential predictor of the resilience metric used by authors, followed by the methodological approach and disturbance pattern (Fig. 4a). Specifically, removing information on these three study attributes causes the largest decline in the model’s predictive performance. Other study attributes contribute comparatively little to the predictive performance of our model. Within the measured variables (Fig. 4b), studies quantifying process-based indicators (*e.g.,* demographic or food-web parameters) are disproportionately associated with rare resilience metrics, defined here as single, uncommon metrics (<5% usage), such as resistance and invariability. We also find that studies that measure quantity-type variables, such as abundance and biomass, are more likely to adopt latitude (*i.e.*, the maximum magnitude of disturbance a system can tolerate before shifting to an alternative stable state^36^) as the resilience metric. In contrast, structural variables, like community composition or diversity, are most frequently associated with recovery degree, whereas functional response variables, such as productivity or respiration rate, most commonly correspond to recovery rate. Methodological approach further differentiates metric selection (Fig. 4c). In particular, latitude is disproportionately adopted in model-based simulations and laboratory experiments relative to field-based approaches. By contrast, field experimental studies most frequently employ recovery degree, whereas recovery rate is used with similar probability across all methodological approaches. Disturbance regime shows a similar structuring effect (Fig. 4d). Studies examining pulse disturbances show higher adoption of recovery degree, recovery time and multidimensional quantification than those focused on press disturbances. Conversely, press-disturbance studies more frequently adopt latitude and recovery rate. Full conditional probability distributions for all predictor categories are provided in Supplementary Table 6. Taken together, these results reveal a field in which resilience is quantified using diverse metrics that are only weakly constrained by study context, yet consistently fall within a shared set of dimensions.

**Figure 4.**
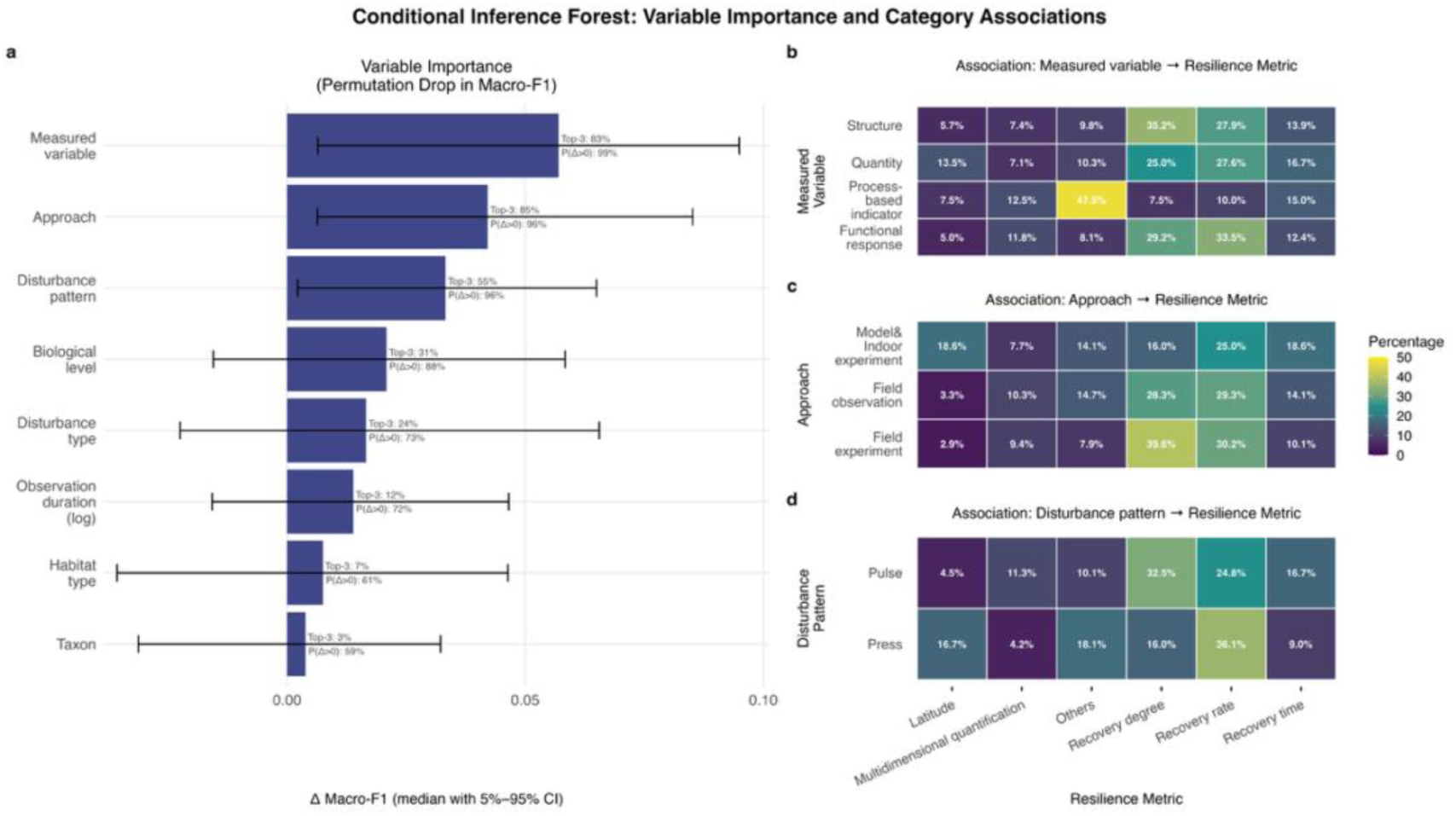
Conditional inference forest analysis of study attributes associated with resilience metrics. (a) Permutation variable importance, measured as the median change in macro-averaged F1 score (Δ macro-F1) when each predictor is permuted across held-out data. The conditional inference forest comprised 2,000 trees with three candidate predictors considered at each split. Error bars indicate the 5–95% interval across repeated stratified cross-validation runs. Labels on the right report how frequently each predictor ranked among the three most important variables and the probability that Δ macro-F1 > 0 across runs. (b–d) Row-normalised heatmaps showing the distribution of resilience metrics across categories of the three most influential predictors: measured variable (b), study approach (c) and disturbance pattern (d). Values indicate row percentages. These panels illustrate how study design and disturbance context are associated with the choice of resilience metrics.

## Discussion

Resilience has long been characterised by conceptual and metric inconsistencies in ecology^9,26^. Here, we use large language models to extract information at scale across 594 publications published between 1977 and 2025 to assess past and current practices in resilience research. We find that the quantification of resilience in ecology remains as diverse in recent publications as in the past, with recovery rate (*sensu* Van Nes & Scheffer^37^) and recovery degree (*sensu* Van Ruijven & Berendse^38^) being the relatively most common metrics. Applying conditional inference forests to test whether inconsistency in resilience quantification relates to study context and author choice, we further find that key attributes of each study such as level of biological organisation or methodological approach show only weak associations with how authors chose to quantify resilience in their systems. This pattern indicates that resilience metrics are not strongly constrained by the inherent characteristics of ecological subfields. Our findings highlight opportunities to reduce communication barriers and foster evidence-based consensus. Below we propose a roadmap to achieve such a synthetic view of resilience in ecology.

### Limited progress in unifying the resilience concept

Recent studies continue to emphasise the persistence of diversity and conceptual fragmentation in resilience quantification^24,39^. Similarly, we find that no single resilience metric has achieved dominance. The two most commonly used metric categories in our dataset are recovery rate and recovery degree, corresponding to the rate of return to equilibrium following disturbance^37^ and the deviation of the post-recovery state from the original stable state^38^, respectively. Notably, recovery degree is not among the two resilience concepts most frequently discussed in the theoretical literature^9,26^, namely ecological resilience (*i.e.,* the maximum disturbance a system can absorb without shifting to an alternative stable state, represented as latitude in this study^15^) and engineering resilience (*i.e.,* the rate of return to equilibrium following perturbation, represented as recovery rate in this study^18^). The prominence of recovery degree as a resilience metric may instead reflect practical and conceptual advantages. Recovery degree can convey information similar to recovery rate when the endpoint of recovery cannot be clearly determined^22^. Moreover, recovery degree accommodates situations in which systems do not fully return to their pre-disturbance state, thereby aligning conceptually with latitude and the possibility of multiple stable^10^. For example, some studies use recovery degree to assess whether regime shifts have occurred^40^. These practical and conceptual connections between recovery degree and the two most widely discussed resilience concepts may help explain why recovery degree emerges as one of the most frequently adopted metrics.

Our findings suggest that past attempts to clarify and unify the concept of resilience in ecology have not yet reduced the diversity of resilience metrics in use^10,26,33^. Nevertheless, more than 90% of our examined studies that quantify resilience using direct metrics (rather than proxies) draw from six common metric categories. This pattern suggests that the multidimensional nature of resilience may be amenable to integration within a unified framework. If such a framework provides a set of dimensions encompassing the principal aspects of resilience studied by ecologists, and gains broad uptake, it could help reduce dialectal fragmentation across subfields. Such a framework would benefit both increasingly emphasised integrative research^5,32^ and the translation of resilience science into conservation practice^13,14^. We contend that previous efforts to unify resilience may have struggled partly because they overlooked recovery degree as a prominent dimension of resilience in practice. For example, Ingrisch and Bahn emphasised recovery rate and resistance as core elements of resilience^33^, and Van Meerbeek et al. advocated framing resilience primarily in terms of recovery rate within a broader stability framework^10^. However, given that recovery degree is among the most frequently used metrics in the journals we examined, such frameworks may encounter challenges in achieving widespread practical adoption.

Two notable trends emerge from our findings. First, multidimensional quantifications of resilience have increased over the past decade, although they remain rare, consistent with previous assessments^14^. This trend may reflect increasing sophistication in ecological monitoring and experimental methodologies, which generate more diverse data types^41,42^. This trend may also indicate growing recognition of the conceptual complexity of resilience and the influence of advocates for multidimensional frameworks^26,33,43^. Second, inference-based studies have increased over the past decade and now constitute approximately 25% of resilience research in ecology. Such studies are often conducted at large spatial scales, where proxies such as land-use types are used to infer resilience^44^. This trend raises concerns regarding the translation of resilience research into conservation and management. In contexts where conceptual definitions remain fragmented, proxies may ambiguously represent different resilience dimensions. Moreover, large-scale studies often exert disproportionate influence on policymaking^45^. Weakly specified proxies may therefore widen the gap between academic discourse and conservation practice^46^.

### Diversity and gaps in study contexts

Our analysis of study attributes reveals substantial diversity, identifiable patterns, and notable gaps within the scope of the journals examined. Different levels of biological organisation tend to favour particular methodological approaches and measured variables, yet nearly all combinations appear somewhere in high-impact ecological journals. This diversity suggests that, with improved comparability, resilience research could generate insights across systems and scales^26,32^

Certain study contexts appear underrepresented and merit further attention. These gaps include population studies that neglect structural population changes when assessing resilience, and the relative scarcity of non-manipulative observational studies of chemical disturbances. In additional, geographic clustering is pronounced within the examined journals. Resilience research is concentrated in North America, Europe, and China, consistent with broader geographic biases documented in ecology^47^. However, these research hotspots do not coincide with regions experiencing the highest disturbance exposure and biodiversity loss^48^. Fire disturbance provides a specific example: sub-Saharan African savannas, Australia and parts of Eastern Europe are global wildfire hotspots^49^, yet wildfire-focused resilience studies from these regions are scarce in our dataset. These disparities may indicate a need for stronger international collaboration targeting biodiversity and disturbance hotspots. They may also reflect structural inequities in research funding and long-term research infrastructure, which influence publication patterns in high-impact journals^50,51^. It is important to note, however, that the gaps identified here reflect only the journals included in our dataset and may not represent the full ecological research landscape.

### Toward a cross-subfield resilience framework

The weak predictive relationship between study context and the choice of resilience metric suggests a high potential for achieving greater consensus across ecological subfields. Our findings suggest that ecologists often treat resilience metrics as a flexible toolkit, selecting metrics suited to specific questions, rather than adhering to rigid conventions imposed by subfield identity. We propose that this flexibility implies that a unified resilience framework could be applied across diverse research contexts and levels of biological organisation, potentially facilitating integrative efforts increasingly emphasised in ecology^5,32^. Certain study-level effects, nevertheless, remain meaningful. For example, latitude—an inherently challenging dimension of resilience to quantify^26,33^—is more frequently employed in model-based simulations and laboratory experiments, and more often examined under press disturbances. This pattern underscores the substantive distinction between systems characterised by single equilibria and those with multiple or no stable equilibria^52^. The latter typically require greater experimental control and longer observational periods for adequate characterisation^17^. A unified framework of resilience in ecology should therefore not be interpreted as requiring individual studies to quantify all dimensions of resilience. Rather, it should enable studies to locate themselves explicitly within a shared conceptual structure, thereby facilitating synthesis, comparison and gap identification.

As resilience research in ecology continues to expand, the coming decade offers an opportunity for ecologists, policy makers, and practitioners to converge on clearer definitions and quantification strategies. In a landscape where no single metric dominates yet several common categories are widely shared, we advocate for a unifying framework encompassing multiple resilience dimensions. Such a framework could reduce terminological conflict^10^ and help the community recognise that differences in metrics reflect distinct resilience components rather than conceptual inconsistencies.

We propose that an effective resilience framework should satisfy two key conditions. First, it should define a set of resilience dimensions that are not inherently correlated and provide example metrics for quantifying each dimension^53^. Second, said resilience framework should accommodate systems lacking a single equilibrium, thereby incorporating dimensions such as recovery degree and latitude. Such a framework would not require every study to address all dimensions. Instead, it would encourage explicit specification of the resilience component being quantified. Researchers could then combine complementary dimensions into composite metrics where appropriate, such as recovery time, which integrates resistance and recovery rate^26,33^. Inference-based studies would likewise be able to clarify which resilience dimensions their proxies represent. Ultimately, clearer specification could extend to policy and conservation practice, reducing ambiguity surrounding resilience and facilitating more explicit objective-setting^13,14^. Under a shared framework, practitioners could identify concrete resilience goals aligned with management needs and more effectively draw upon ecological evidence.

## Methods

### Literature search

To examine how ecologists have defined and quantified resilience over time, we conducted a systematic synthesis following PRISMA-EcoEvo guidelines^54^. The search, screening and inclusion process is summarised in Supplementary Fig. 5. On the 14th of January 2026, we searched Scopus and the Web of Science Core Collection using the query: ‘resilience’ AND (‘disturbance’ OR ‘perturbation’ OR ‘stress*’ OR ‘disruption’ OR ‘shift’ OR ‘stability’ OR ‘resistance’ OR ‘recovery’ OR ‘elasticity’ OR ‘compensation’). These terms are commonly used in disturbance ecology (*e.g*., ^43,55–57^). We required at least one of these terms to appear in the title, abstract, or author-provided keywords to ensure that resilience was examined within a relevant ecological context. To ensure disciplinary relevance and capture influential contributions to resilience research in ecology, we restricted our search to original research articles published in 15 selected journals: Nature, Science, Proceedings of the National Academy of Sciences, and top 12 ecology journals ranked by the Google Scholar h5-index (Supplementary Table 7). In addition, we only included articles published before the 1st of January to avoid the need for continual updates to the search results. These criteria resulted in a dataset of 1,959 peer-reviewed articles spanning from 1977 to 2025 after removing duplicates.

### Study inclusion and data collection

To reduce the number of full papers that needed to be downloaded and filtering, we conducted screening in two stages: abstract screening followed by full-text screening. Our screening retained studies that: (1) quantify resilience in an ecological context, and (2) are not exclusively focused on abiotic response variables. Of the 1,959 articles, 1,227 (63%) passed abstract screening and 594 (30%) passed full-text screening.

To examine how ecologists quantify resilience and the contexts in which they apply it, we extracted structured data from the 594 articles that met our inclusion criteria. From each article, we recorded a set of study attributes, including: (1) Resilience quantification (see Table 1), *i.e.,* how resilience is quantified; (2) Measured variable type for resilience quantification (see Supplementary Table 1); (3) Level of biological organisation (*i.e.,* individual, population, community, ecosystem, and landscape); (4) Approach (*i.e*., modelling, laboratory experiment, field experiment, and/or field observation); (5) Disturbance type (*e.g*., drought, fire; see Supplementary Table 8); (6) Disturbance pattern, categorised as pulse (short-term discrete events) or press (long-term continuous pressures)^34^; (7) Observation duration, *i.e.,* the interval between the first and last observations; (8) Habitat type (as per IUCN^35^); (9) Taxonomic group; In addition, we also recorded other study attributes such as the conceptual framework of the study (see Supplementary Table 9), study location, the country of the first author’s institution, and the approach to examine disturbances (*i.e.,* model, manipulated, unmanipulated) (Supplementary Methods).

To facilitate the collection of these data, we classified the studies into direct-response vs. inference-based studies. We based this distinction on how resilience was measured: direct-response studies quantify resilience from measurements taken directly on the system’s response to disturbance^58^. Inference-based studies infer resilience from indicators or models that act as proxies rather than direct response measurements^59^. We coded disturbance-related variables (*e.g*., disturbance type, observation duration) for direct-response studies only, as such information was typically unavailable or ambiguous in inference-based studies.

### Large language model

To reduce the time and labour required to process more than 2,000 collected publications, we used large language models (LLMs) to assist with abstract screening, full-text screening, and data extraction. We implemented all LLM-related tasks in Python 3.12.7 via OpenAI’s application programming interface. We used OpenAI’s GPT-5.2 for all tasks, with the temperature fixed at 0. We provided abstracts to the LLM via CSV files and full-text content by article PDFs with the PyPDF2 package. For transparency and reproducibility, we maintained a spreadsheet linking each output file to the exact prompt text and model. For full methodological details of the LLM-assisted screening and information extraction, see ^60^.

To keep the workflow tractable and reproducible, we framed each task as a well-specified prompt to the model, comprising multiple structured questions. In screening, each question asked whether a particular inclusion criterion was met. In data extraction, each question targeted a predefined extraction variable. When submitting prompts to the model for each task, we included the abstract or full text of only one article at a time to avoid any cross-article influence on the outputs. Prompts requested a compact JSON reply with named fields aligned to our variables. We parsed the JSON into a table and applied minimal cleaning, such as letter case harmonising (e.g., “Forest” → “forest”).

To develop and refine the prompts, we first drafted an initial version for each task, and then tested these prompts using LLMs on a subset of articles with human-coded answers and measured their performance against human decisions. After examining common errors in the LLM’s outputs (e.g., without explicit instruction, the model may classify all studies using remote-sensing data as inference-based, assuming that such data do not directly observe the system), we revised the prompts based on the observed errors and, where useful, brief consultations with LLMs. Finally, we re-tested the prompts to assess whether performance improved and observed errors were reduced. Following Moorthy et al.^60^, when further prompt revisions yielded only marginal improvements in performance, we validated the final few versions on a separate subset of articles. The performance of each prompt had to meet our predefined acceptance thresholds (see the following paragraph) to pass validation. We then selected the prompt with the best performance from validation results for deployment (i.e., apply to all the remaining articles). After each deployment, we audited a random sample of 50 articles from the outputs to confirm that performance remained above our acceptance thresholds^60^.

To implement this develop–validate–deploy procedure, we used task-specific sample sizes and acceptance performance thresholds. For abstract screening, we developed the prompt on 50 articles, validated on 100, and then deployed on the rest of 1,809 articles. We accepted a training–validation set comprising less than 10% of the full sample because the abstract-screening stage served as a coarse pre-filter. Our primary objective at this stage was to reduce false negatives; accordingly, while we only require ≥ 80% consistency with human judgments, we set a strict performance threshold of ≤ 2% false negatives. For full-text screening, we developed the prompt on 100 articles, validated on 100, and deployed on the rest of 1,027. The performance thresholds at this stage are ≥ 85% consistency with human and Cohen’s κ ≥ 0.7. Cohen’s κ measures inter-rater reliability beyond chance agreement^61^. A κ value ≥ 0.7 lies in the “substantial agreement” range (κ = 0.61–0.80)^62^. For data extraction, we developed the prompt on 75 articles, validated on 75, and deployed the prompt to 444 papers. The performance threshold is F1-score ≥ 0.80 for every extraction variable. F1-score is an accuracy metric^63^. F1-score ≥ 0.80 is generally considered to indicate strong model performance in classification tasks^64^.

Although all extraction variables met the predefined F1-score thresholds after the data extraction using LLMs, we identified a systematic bias in the extraction of the response variable in our analyses, resilience quantification. Specifically, the model tended to return a single resilience quantification even when a study explicitly adopted a resilience framework in which resilience was defined as comprising multiple quantitative components. While some studies classified under a resilience framework (see Supplementary Table 9) acknowledge the multidimensional nature of resilience but nevertheless operationalise it using a single quantification^65–67^, the proportion of such cases in the model outputs was markedly higher than expected based on comparisons with manually extracted data. Because quantification constituted a central variable in our analyses, we manually reviewed all direct-response studies classified under a resilience framework but assigned a single quantification value by the model (n = 65). This review resulted in the addition of systematically omitted quantification values for 25 studies. Full details of all LLM-related tasks are provided in Supplementary Methods.

### Statistical analyses

To test whether study contexts predict how ecologists quantify resilience, we trained a conditional inference forest to predict the resilience quantification using eight study attributes as predictors. Traditional random forests are prone to variable selection bias^68^. The conditional inference forest avoids this issue by selecting splitting variables through permutation-based statistical tests. To do so, we used the party R package^69^. We optimised hyperparameters through stratified cross-validation: number of variables considered at each split = 3, additional significance threshold = 0, and 2,500 trees. Full details of model specification, hyperparameter tuning, sensitivity analyses, complementary analyses, and permutation-based importance results are provided in Supplementary Methods and Results.

To avoid our response variable showing class imbalance, where the most frequent category exceeds the least frequent by more than 4:1^70^, we reclassified the data into six categories: recovery rate, recovery degree, recovery time, multidimensional quantification, latitude and others. All samples with more than one resilience metric fall under multidimensional quantification. All remaining metric types with frequencies below one quarter of the largest class (*i.e.*, 33 samples) are grouped under others. This reclassification resulted in a final class ratio of 3.46:1 between the majority class (recovery rate) and the minority (latitude). To mitigate the potential impact of unequal class frequencies, we employed stratified cross-validation to ensure proportional class representation across all folds^71^ and used macro-averaged F1 score as the primary performance metric, which treats all classes equally regardless of size^72^.

We included a comprehensive set of predictors because several study-context factors likely drive the diversity in current resilience-quantification practices. These predictors include variables (2) through (9) described in Methods (also see Supplementary Methods). Before model fitting, we excluded all articles classified as inference-based because the disturbance-related predictors required by our models were unavailable for these studies. To account for potential differences in thematic scope among journals and potential influence from geographical academic networks, we also ran a complementary analysis: we included the journal in which a paper was published and the country of the first author’s institution as additional predictors. For all predictors, we grouped categories with fewer than 20 samples to reduce data sparsity.

To assess the contribution of each predictor to model performance, we used permutation-based importance analysis after the model fitting. This analysis helped us to understand how each study-context factor influenced researchers’ choice of resilience metric. We did so by applying 30 repeats of five-fold cross-validation^73^ and evaluating model performance using the strict macro-F1-score^74^. This metric averages F1-scores across all categories with equal weight, regardless of class size. Finally, we calculated group-wise permutation importance by jointly permuting predefined predictors and measuring the associated drop in macro-F1. This approach allowed us to rank predictors according to their relative importance^75^.

## Data Availability

The study-level attributes extracted from the analysed publications are provided in Supplementary Data 3. These include both manually curated variables and variables extracted using large language models, and comprise the dataset used for all analyses.

## Code Availability

R scripts used for data curation and statistical analyses are available at GitHub (https://github.com/homewaylin666/Systematic-Review-of-Resilience.git).

## Notes

https://github.com/homewaylin666/Systematic-Review-of-Resilience.git

### Competing Interest Statement

The authors have declared no competing interest.

