## Supplementary Information for "Still Lost in Definitions: How ‘Resilience’ Is Used in Ecology"

#### Contents

**S1. Supplementary Tables** **2**

Supplementary Table 1 2

Supplementary Table 2 4

Supplementary Table 3 7

Supplementary Table 4 9

Supplementary Table 5 11

Supplementary Table 6 12

Supplementary Table 7 18

Supplementary Table 8 19

Supplementary Table 9 20

**S2. Supplementary Figures** **21**

Supplementary Fig. 1 21

Supplementary Fig. 2 22

Supplementary Fig. 3 23

Supplementary Fig. 4 24

Supplementary Fig. 5 25

**S3. Supplementary Methods** **26**

Details of data collection 26

Details of LLM-related task 27

Details of conditional inference forest 30

**S4. Supplementary Results** **32**

Sensitivity tests of conditional inference forest 32

An alternative conditional inference forest for exploration 34

Results from half of the journals 36

**S5. Supplementary Information Reference** **42**

### S1 Supplementary Tables

**Supplementary Table 1.** Classification of measured variables used to quantify resilience. A measured variable is the measured quantity whose response relative to disturbance is used to quantify resilience. Variables are grouped into five types: Quantity, Structure, Process-based indicators, Functional response, and Environmental context. Within each category, we recognise subtypes (e.g., within Structure: community composition, age structure, and alpha diversity). Examples are illustrative rather than exhaustive. Studies may report multiple measured variables. The dataset codes every variable used to quantify resilience, so a single study can fall into multiple subtypes. This classification underpins subsequent analyses of associations among organisational level, methodological approach, and measured variable category, and it informs the conditional inference forest analysis.

| Measured variable type | Subtype | Examples |
| --- | --- | --- |
| Quantity | Cover | plant cover, coral cover, *etc.* |
| | Density | seed density, coral density *etc.* |
| | Abundance | population abundance, *etc.* |
| | Biomass | plant biomass, fish biomass, *etc.* |
| Structure | Composition | community composition, beta diversity, relative frequencies or ratios of specified species., *etc.* |
| | | age structure, such as relative density of different life history stages |
| | Diversity | alpha diversity, such as richness, evenness, and not limited to species but also including genetic diversity, functional diversity, *etc.* |
| Process-based indicator | Demography parameter | survival rate, recruitment rate, mortality rate, growth rate, *etc.* |
| | Interaction | species interaction proxies, such as seed dispersal and prey capture |
| | | food-web parameters, such as energy flows between trophic levels, trophic links, the domain of attraction of a food-web |
| | | other interaction-network parameters, such as modularity, nestedness, βeff |

| | | |
|---|---|---|
| Functional response | Ecosystem function | productivity, chlorophyll a concentration, biomass as the proxy of productivity, vegetation indices (e.g. NDVI) as proxies of productivity etc. |
| | | water storage, water transparency, *etc.* |
| | | biogeochemical or energy output |
| | Physiological indicator | metabolism, respiration rate, protein synthesis rate, morphology, N ratio, gene expression patterns, greenness *etc.* |
| | Growth | individual growth, such as tree-ring growth, basal area increment, height, *etc.* |
| Environmental context | Abiotic parameter | temperature, moisture, *etc.* |
| | Land use | land use type |
| | Regional network parameter | network efficiency; network robustness; largest connectivity subgraph, *etc.* |
| | Socio-eco parameter | human population density, Industrial wastewater emissions, social electricity consumption *etc.* |

**Supplementary Table 2. The research contexts of resilience-quantified studies exhibit non-random association.** Results of chi-square analyses examining pairwise associations among study attributes (corresponding to Fig. 3). Each row represents a cell in the contingency table, showing the observed count, expected frequency under independence, standardised residual (z-score), and the direction of deviation. Grey-shaded cells denote significant deviations ($|z| \geq 1.96$, $p < 0.05$). Positive z-scores indicate an excess of studies relative to expectation, whereas negative z-scores indicate a deficit. Direction is coded as EXCESS (standardised residual > 1.96), DEFICIT (standardised residual < −1.96) or NS (not significant; $|z| \leq 1.96$). Abbreviations: MD = model-based simulation; LM = indoor (laboratory) experiment; FM = field experiment; FO = field observation. Pairwise comparisons include: (1) Level *vs.* Approach, linking organisational level (Landscape, Ecosystem, Community, Population, Individual) with research approach; (2) Level *vs.* Measured variable, linking organisational level with the focal variable type (Environmental Context, Functional Response, Structure, Process-based Indicator, Quantity); and (3) Approach *vs.* Measured variable, linking research approach with the focal variable type.

| Pair | Variable1 | Variable2 | count | expected | z-score | direction |
|---|---|---|---|---|---|---|
| Level *vs.* Approach | Landscape | MD | 23 | 12.641 | 3.501 | EXCESS |
| | Ecosystem | MD | 50 | 50.801 | -0.161 | NS |
| | Community | MD | 38 | 48.893 | -2.213 | DEFICIT |
| | Population | MD | 28 | 19.557 | 2.359 | EXCESS |
| | Individual | MD | 1 | 8.109 | -2.948 | DEFICIT |
| | Landscape | LM | 0 | 1.535 | -1.318 | NS |
| | Ecosystem | LM | 6 | 6.169 | -0.086 | NS |
| | Community | LM | 4 | 5.937 | -1.000 | NS |
| | Population | LM | 5 | 2.375 | 1.864 | NS |
| | Individual | LM | 2 | 0.985 | 1.070 | NS |
| | Landscape | FM | 1 | 12.460 | -3.892 | DEFICIT |
| | Ecosystem | FM | 45 | 50.075 | -1.027 | NS |
| | Community | FM | 64 | 48.194 | 3.227 | EXCESS |
| | Population | FM | 17 | 19.278 | -0.640 | NS |
| | Individual | FM | 11 | 7.993 | 1.253 | NS |
| | Landscape | FO | 29 | 26.365 | 0.759 | NS |
| | Ecosystem | FO | 112 | 105.956 | 1.038 | NS |
| | Community | FO | 99 | 101.976 | -0.515 | NS |
| | Population | FO | 32 | 40.791 | -2.093 | DEFICIT |
| | Individual | FO | 20 | 16.913 | 1.091 | NS |
| | Landscape | Environmental context | 37 | 6.150 | 13.873 | EXCESS |
| | Ecosystem | Environmental context | 30 | 24.717 | 1.417 | NS |

| | | | | | | |
|---|---|---|---|---|---|---|
| | Community | Environmental context | 1 | 23.672 | -6.139 | DEFICIT |
| | Population | Environmental context | 0 | 9.515 | -3.538 | DEFICIT |
| | Individual | Environmental context | 0 | 3.945 | -2.177 | DEFICIT |
| | Landscape | Functional response | 3 | 15.195 | -3.884 | DEFICIT |
| | Ecosystem | Functional response | 117 | 61.065 | 10.623 | EXCESS |
| | Community | Functional response | 8 | 58.485 | -9.681 | DEFICIT |
| | Population | Functional response | 13 | 23.509 | -2.767 | DEFICIT |
| | Individual | Functional response | 27 | 9.747 | 6.741 | EXCESS |
| | Landscape | Structure | 3 | 9.587 | -2.465 | DEFICIT |
| | Ecosystem | Structure | 7 | 38.529 | -7.035 | DEFICIT |
| Level *vs.* Measured variable | Community | Structure | 96 | 36.901 | 13.314 | EXCESS |
| | Population | Structure | 0 | 14.833 | -4.589 | DEFICIT |
| | Individual | Structure | 0 | 6.150 | -2.823 | DEFICIT |
| | Landscape | Process-based indicator | 5 | 4.432 | 0.296 | NS |
| | Ecosystem | Process-based indicator | 7 | 17.811 | -3.354 | DEFICIT |
| | Community | Process-based indicator | 22 | 17.058 | 1.548 | NS |
| | Population | Process-based indicator | 13 | 6.857 | 2.643 | EXCESS |
| | Individual | Process-based indicator | 2 | 2.843 | -0.538 | NS |
| | Landscape | Quantity | 5 | 17.637 | -3.863 | DEFICIT |
| | Ecosystem | Quantity | 52 | 70.879 | -3.441 | DEFICIT |
| | Community | Quantity | 77 | 67.884 | 1.678 | NS |
| | Population | Quantity | 56 | 27.287 | 7.256 | EXCESS |
| | Individual | Quantity | 5 | 11.314 | -2.368 | DEFICIT |
| | MD | Environmental context | 24 | 16.130 | 2.387 | EXCESS |
| | LM | Environmental context | 1 | 1.973 | -0.748 | NS |
| | FM | Environmental context | 4 | 16.014 | -3.652 | DEFICIT |
| | FO | Environmental context | 39 | 33.884 | 1.320 | NS |
| | MD | Functional response | 21 | 39.850 | -4.048 | DEFICIT |
| | LM | Functional response | 7 | 4.874 | 1.157 | NS |
| | FM | Functional response | 39 | 39.563 | -0.121 | NS |
| Approach *vs.* Measured variable | FO | Functional response | 101 | 83.713 | 3.158 | EXCESS |
| | MD | Structure | 9 | 25.143 | -4.073 | DEFICIT |
| | LM | Structure | 1 | 3.075 | -1.327 | NS |
| | FM | Structure | 30 | 24.963 | 1.274 | NS |
| | FO | Structure | 66 | 52.819 | 2.829 | EXCESS |
| | MD | Process-based indicator | 17 | 11.623 | 1.887 | NS |
| | LM | Process-based indicator | 1 | 1.422 | -0.375 | NS |
| | FM | Process-based indicator | 4 | 11.539 | -2.652 | DEFICIT |
| | FO | Process-based indicator | 27 | 24.416 | 0.771 | NS |
| | MD | Quantity | 68 | 46.254 | 4.482 | EXCESS |

| | | | | | |
|---|---|---|---|---|---|
| LM | Quantity | 7 | 5.6570 | 0.7015 | NS |
| FM | Quantity | 61 | 45.9215 | 3.1154 | EXCESS |
| FO | Quantity | 59 | 97.1672 | -6.6922 | DEFICIT |

**Supplementary Table 3. Press disturbances were significantly less common than pulse disturbances within manipulated disturbances.** Results of chi-square analyses examining pairwise associations among disturbance manipulation, disturbance type, and disturbance pattern (corresponding to Extend Data Fig. 1a–c). Each row represents a cell in the contingency table, showing the observed study count, the standardised residual (z-score), and the direction of deviation relative to independence. Grey-shaded cells denote significant deviations ($|z| \geq 1.96$, $p < 0.05$). Positive z-scores indicate an excess of studies relative to expectation, whereas negative z-scores indicate a deficit. Direction is coded as EXCESS (standardised residual > 1.96), DEFICIT (standardised residual < −1.96) or NS (not significant; $|z| \leq 1.96$). Abbreviations: Manip. = manipulated; Unmanip. = unmanipulated; LID = Land Use and Infrastructure Development; BRU = Biological Resource Use. Patterns: Press = sustained or chronic disturbance as treated in the study; Pulse = discrete event(s) as treated in the study.

| Pair | Variable1 | Variable2 | count | z-score | direction |
|---|---|---|---|---|---|
| Disturbance manipulation *vs.* Disturbance type | Model | Fire | 7 | -1.376 | NS |
| | Manip. | Fire | 11 | -1.845 | NS |
| | Unmanip. | Fire | 41 | 2.746 | EXCESS |
| | Model | Climatic | 36 | -1.699 | NS |
| | Manip. | Climatic | 46 | -4.200 | DEFICIT |
| | Unmanip. | Climatic | 156 | 5.138 | EXCESS |
| | Model | Physical | 0 | -1.267 | NS |
| | Manip. | Physical | 0 | -1.701 | NS |
| | Unmanip. | Physical | 7 | 2.530 | EXCESS |
| | Model | Physical | 4 | -0.895 | NS |
| | Manip. | Physical | 8 | -0.514 | NS |
| | Unmanip. | Physical | 20 | 1.162 | NS |
| | Model | Chemical | 5 | -1.322 | NS |
| | Manip. | Chemical | 31 | 6.122 | EXCESS |
| | Unmanip. | Chemical | 9 | -4.536 | DEFICIT |
| | Model | Resource | 9 | 2.761 | EXCESS |
| | Manip. | Resource | 12 | 2.687 | EXCESS |
| | Unmanip. | Resource | 1 | -4.588 | DEFICIT |
| | Model | Biotic | 10 | -1.427 | NS |
| | Manip. | Biotic | 26 | 0.818 | NS |
| | Unmanip. | Biotic | 43 | 0.366 | NS |
| | Model | LID | 7 | 0.416 | NS |
| | Manip. | LID | 0 | -3.775 | DEFICIT |
| | Unmanip. | LID | 26 | 3.107 | EXCESS |
| | Model | BRU | 13 | 3.315 | EXCESS |

| | | | | | |
|---|---|---|---|---|---|
| | Manip. | BRU | 8 | -0.514 | NS |
| | Unmanip. | BRU | 11 | -2.110 | DEFICIT |
| | Model | Structural | 23 | 3.293 | EXCESS |
| | Manip. | Structural | 37 | 4.669 | EXCESS |
| | Unmanip. | Structural | 10 | -6.802 | DEFICIT |
| Disturbance manipulation *vs.* Disturbance pattern | Model | Press | 39 | 2.989 | EXCESS |
| | Manip. | Press | 36 | -2.620 | DEFICIT |
| | Unmanip. | Press | 81 | 0.156 | NS |
| | Model | Pulse | 48 | -2.989 | DEFICIT |
| | Manip. | Pulse | 119 | 2.620 | EXCESS |
| | Unmanip. | Pulse | 175 | -0.156 | NS |
| Disturbance type *vs.* Disturbance pattern | Fire | Press | 8 | -3.418 | DEFICIT |
| | Climatic | Press | 83 | 0.579 | NS |
| | Geophysical | Press | 1 | -1.062 | NS |
| | Hydrological | Press | 8 | -1.107 | NS |
| | Chemical | Press | 16 | 0.257 | NS |
| | Resource | Press | 14 | 2.888 | EXCESS |
| | Biotic | Press | 33 | 1.855 | NS |
| | LID | Press | 16 | 1.935 | NS |
| | BRU | Press | 17 | 2.476 | EXCESS |
| | Structural | Press | 10 | -3.545 | DEFICIT |
| | Fire | Pulse | 52 | 3.418 | EXCESS |
| | Climatic | Pulse | 158 | -0.579 | NS |
| | Geophysical | Pulse | 6 | 1.062 | NS |
| | Hydrological | Pulse | 25 | 1.107 | NS |
| | Chemical | Pulse | 30 | -0.257 | NS |
| | Resource | Pulse | 9 | -2.888 | DEFICIT |
| | Biotic | Pulse | 45 | -1.855 | NS |
| | LID | Pulse | 17 | -1.935 | NS |
| | BRU | Pulse | 15 | -2.476 | DEFICIT |
| | Structural | Pulse | 60 | 3.545 | EXCESS |

**Supplementary Table 4. The duration of the study varies significantly across the disturbance types examined.** Results of one-way ANOVA and Tukey's HSD post-hoc tests examining variation in observation duration ($\log_{10}$ days) across disturbance categories (corresponding to Fig. 4d–f). Each ANOVA tested differences in mean observation duration among factor levels of disturbance manipulation, disturbance type, or disturbance pattern. Significant ANOVA results were followed by post-hoc Tukey's HSD pairwise comparisons between levels of each factor. The upper section lists ANOVA results with the F-statistic, numerator and residual degrees of freedom, and p-values. The lower section lists significant Tukey's HSD contrasts ($p < 0.05$) with estimated mean differences (estimate) and 95 % confidence-interval limits. Positive estimates indicate higher mean $\log_{10}$ observation durations in the first group relative to the second. Abbreviations: Manip. = manipulated; Unmanip. = unmanipulated; LID = Land Use and Infrastructure Development; BRU = Biological Resource Use; Press = sustained or chronic disturbance as treated in the study; Pulse = discrete event(s) as treated in the study.

| Test Type | Factor | term | statistic | df | Residuals df | p value |
|---|---|---|---|---|---|---|
| ANOVA | Manipulation *vs.* Duration | Disturbance manipulation | 87.608 | 2 | 481 | <0.001 |
| | Type *vs.* Duration | Disturbance type | 10.540 | 9 | 591 | <0.001 |
| | Pattern *vs.* Duration | Disturbance pattern | 10.664 | 1 | 466 | 0.001 |
| Test Type | Factor | term | estimate | conf low | conf high | p value |
| Tukey HSD | Disturbance manipulation | Manip.-Model | -3.226 | -3.965 | -2.487 | <0.001 |
| | | Unmanip.-Manip. | 2.944 | 2.379 | 3.509 | <0.001 |
| | Disturbance type | Chemical-Fire | -2.661 | -4.322 | -1.000 | <0.001 |
| | | Resource-Fire | -3.366 | -5.468 | -1.263 | <0.001 |
| | | Structural-Fire | -1.913 | -3.379 | -0.447 | 0.002 |
| | | Chemical-Climatic | -1.947 | -3.318 | -0.577 | <0.001 |
| | | Resource-Climatic | -2.652 | -4.533 | -0.771 | <0.001 |
| | | LID-Climatic | 1.942 | 0.385 | 3.499 | 0.003 |
| | | Structural-Climatic | -1.199 | -2.325 | -0.073 | 0.026 |
| | | Chemical-Geophysical | -4.715 | -8.080 | -1.351 | <0.001 |
| | | Resource-Geophysical | -5.420 | -9.023 | -1.817 | <0.001 |
| | | Structural-Geophysical | -3.967 | -7.240 | -0.695 | 0.005 |
| | | LID-Hydrological | 2.545 | 0.464 | 4.625 | 0.004 |
| | | LID-Chemical | 3.889 | 1.962 | 5.817 | <0.001 |
| | | BRU-Chemical | 2.852 | 0.925 | 4.779 | <0.001 |
| | | Biotic-Resource | 2.243 | 0.205 | 4.281 | 0.018 |
| | | LID-Resource | 4.594 | 2.275 | 6.912 | <0.001 |

| | | | | | | |
|---|---|---|---|---|---|---|
| | | BRU-Resource | 3.557 | 1.238 | 5.875 | <0.001 |
| | | LID-Biotic | 2.351 | 0.608 | 4.094 | <0.001 |
| | | Structural-LID | -3.141 | -4.903 | -1.379 | <0.001 |
| | | Structural-BRU | -2.104 | -3.865 | -0.342 | 0.006 |
| | Disturbance pattern | Pulse-Press | -0.886 | -1.418 | -0.353 | 0.001 |

**Supplementary Table 5. The prediction accuracy of our model varies across resilience metric categories.** Precision, recall, and F1 scores for each resilience metric category. Columns: class label; F1 = harmonic mean of precision and recall; TP = true positives; FP = false positives; FN = false negatives. These per-class metrics complement the aggregate macro-F1 and indicate which classes are reliably identified versus systematically confused. All studies with more than one resilience metric fall under multidimensional quantification. Others include all remaining metric types, such as resistance, invariability, *etc.*

| Metric category | Precision | Recall | F1 | TP | FP | FN |
|---|---|---|---|---|---|---|
| Latitude | 0.454 | 0.224 | 0.300 | 262 | 315 | 908 |
| Multidimensional quantification | 0.196 | 0.016 | 0.029 | 21 | 86 | 1299 |
| Others | 0.444 | 0.289 | 0.350 | 520 | 651 | 1280 |
| Recovery degree | 0.402 | 0.598 | 0.481 | 2369 | 3529 | 1591 |
| Recovery rate | 0.366 | 0.505 | 0.425 | 2047 | 3542 | 2003 |
| Recovery time | 0.296 | 0.147 | 0.196 | 304 | 724 | 1766 |

**Supplementary Table 6. Predictor categories exhibit systematic differences in their preferred resilience metrics.** Distribution of resilience metrics across categories of each categorical predictor variable in the conditional inference forest. Percentage = proportion within the predictor category. The number under category is the total sample size for that predictor category These data reveal systematic patterns in resilience metric selection across study contexts (e.g., pulse disturbances favour recovery degree [32.5%] while press disturbances Favor recovery rate [36.1%]). Observation duration (continuous predictor) is excluded as categorical associations are not applicable to continuous variables; its predictive contribution is shown in variable importance analysis (Fig. 4). All studies with more than one resilience metric fall under multidimensional quantification. Others include all remaining metric types, such as invariability, latitude, *etc.* Definitions of categories are provided in the Methods and Supplementary Methods.

| Predictor | Category | Resilience metric | Percentage |
|---|---|---|---|
| Disturbance pattern | Press (n=144) | Recovery rate | 36.11 |
| | | Others | 18.06 |
| | | Latitude | 16.67 |
| | | Recovery degree | 15.97 |
| | | Recovery time | 9.03 |
| | | Multidimensional quantification | 4.17 |
| | Pulse (n=335) | Recovery degree | 32.54 |
| | | Recovery rate | 24.78 |
| | | Recovery time | 16.72 |
| | | Multidimensional quantification | 11.34 |
| | | Others | 10.15 |
| | | Latitude | 4.48 |
| Measured variable type | Functional response (n=161) | Recovery rate | 33.54 |
| | | Recovery degree | 29.19 |
| | | Recovery time | 12.42 |
| | | Multidimensional quantification | 11.80 |
| | | Others | 8.07 |
| | | Latitude | 4.97 |
| | Process-based indicator | Others | 47.50 |
| | | Recovery time | 15.00 |

| | | | |
|---|---|---|---|
| | (n=40) | Multidimensional quantification | 12.50 |
| | | Recovery rate | 10.00 |
| | | Latitude | 7.50 |
| | | Recovery degree | 7.50 |
| | Quantity (n=156) | Recovery rate | 27.56 |
| | | Recovery degree | 25.00 |
| | | Recovery time | 16.67 |
| | | Latitude | 13.46 |
| | | Others | 10.26 |
| | | Multidimensional quantification | 7.05 |
| | Structure (n=122) | Recovery degree | 35.25 |
| | | Recovery rate | 27.87 |
| | | Recovery time | 13.93 |
| | | Others | 9.84 |
| | | Multidimensional quantification | 7.38 |
| | | Latitude | 5.74 |
| Approach | Field experiment (n=139) | Recovery degree | 39.57 |
| | | Recovery rate | 30.22 |
| | | Recovery time | 10.07 |
| | | Multidimensional quantification | 9.35 |
| | | Others | 7.91 |
| | | Latitude | 2.88 |
| | Field observation (n=184) | Recovery rate | 29.35 |
| | | Recovery degree | 28.26 |
| | | Others | 14.67 |
| | | Recovery time | 14.13 |
| | | Multidimensional quantification | 10.33 |
| | | Latitude | 3.26 |
| | Model-based simulation / Laboratory experiment | Recovery rate | 25.00 |
| | | Latitude | 18.59 |
| | | Recovery time | 18.59 |
| | | Recovery degree | 16.03 |

| | | | |
|---|---|---|---|
| | (n=156) | Others | 14.10 |
| | | Multidimensional quantification | 7.69 |
| Level | Community (n=178) | Recovery degree | 29.78 |
| | | Recovery rate | 27.53 |
| | | Recovery time | 14.61 |
| | | Others | 12.36 |
| | | Multidimensional quantification | 9.55 |
| | | Latitude | 6.18 |
| | Ecosystem / Landscape (n=190) | Recovery rate | 33.16 |
| | | Recovery degree | 24.21 |
| | | Latitude | 12.63 |
| | | Others | 12.63 |
| | | Recovery time | 12.11 |
| | | Multidimensional quantification | 5.26 |
| | Individual (n=33) | Recovery degree | 42.42 |
| | | Multidimensional quantification | 24.24 |
| | | Recovery time | 15.15 |
| | | Recovery rate | 12.12 |
| | | Others | 6.06 |
| | | Latitude | 0.00 |
| | Population (n=78) | Recovery degree | 24.36 |
| | | Recovery rate | 24.36 |
| | | Recovery time | 19.23 |
| | | Others | 15.38 |
| | | Multidimensional quantification | 11.54 |
| | | Latitude | 5.13 |
| Disturbance type | Biotic pressure (n=88) | Recovery degree | 28.41 |
| | | Recovery rate | 18.18 |
| | | Recovery time | 18.18 |
| | | Others | 17.05 |
| | | Latitude | 13.64 |
| | | Multidimensional quantification | 4.55 |

| | | | |
|---|---|---|---|
| | Chemical / Resource (n=50) | Recovery rate | 34.00 |
| | | Recovery degree | 20.00 |
| | | Recovery time | 16.00 |
| | | Multidimensional quantification | 12.00 |
| | | Others | 10.00 |
| | | Latitude | 8.00 |
| | Climatic (n=173) | Recovery degree | 33.53 |
| | | Recovery rate | 30.06 |
| | | Others | 11.56 |
| | | Multidimensional quantification | 10.98 |
| | | Latitude | 6.94 |
| | | Recovery time | 6.94 |
| | Fire (n=55) | Recovery degree | 34.55 |
| | | Recovery rate | 21.82 |
| | | Recovery time | 21.82 |
| | | Others | 10.91 |
| | | Latitude | 5.45 |
| | | Multidimensional quantification | 5.45 |
| | Physical template / Land use change (n=59) | Recovery rate | 37.29 |
| | | Recovery degree | 16.95 |
| | | Recovery time | 15.25 |
| | | Others | 13.56 |
| | | Multidimensional quantification | 10.17 |
| | | Latitude | 6.78 |
| | Structural (n=54) | Recovery rate | 29.63 |
| | | Recovery time | 22.22 |
| | | Recovery degree | 18.52 |
| | | Multidimensional quantification | 11.11 |
| | | Others | 11.11 |
| | | Latitude | 7.41 |
| Taxon | Animal / Animal + Plant | Recovery rate | 29.09 |
| | | Recovery degree | 19.09 |

| | | | |
|---|---|---|---|
| | (n=110) | Others | 18.18 |
| | | Recovery time | 16.36 |
| | | Multidimensional quantification | 10.00 |
| | | Latitude | 7.27 |
| | Microbe included (n=59) | Recovery degree | 33.90 |
| | | Recovery rate | 27.12 |
| | | Recovery time | 18.64 |
| | | Latitude | 8.47 |
| | | Multidimensional quantification | 6.78 |
| | | Others | 5.08 |
| | No specific (n=55) | Recovery rate | 36.36 |
| | | Latitude | 20.00 |
| | | Recovery time | 18.18 |
| | | Others | 14.55 |
| | | Recovery degree | 9.09 |
| | | Multidimensional quantification | 1.82 |
| | Plantae (n=255) | Recovery degree | 33.73 |
| | | Recovery rate | 26.27 |
| | | Recovery time | 11.76 |
| | | Others | 11.37 |
| | | Multidimensional quantification | 10.98 |
| | | Latitude | 5.88 |
| Habitat type | Forest (n=125) | Recovery degree | 36.00 |
| | | Recovery rate | 28.00 |
| | | Others | 16.00 |
| | | Recovery time | 8.00 |
| | | Multidimensional quantification | 7.20 |
| | | Latitude | 4.80 |
| | Marine / Coastal (n=103) | Recovery degree | 23.30 |
| | | Recovery rate | 22.33 |
| | | Recovery time | 22.33 |
| | | Others | 11.65 |

| | | | |
|---|---|---|---|
| | | Multidimensional quantification | 10.68 |
| | | Latitude | 9.71 |
| | No specific / No major (n=57) | Recovery rate | 38.60 |
| | | Latitude | 14.04 |
| | | Recovery time | 14.04 |
| | | Others | 12.28 |
| | | Recovery degree | 12.28 |
| | | Multidimensional quantification | 8.77 |
| | Open barren (n=44) | Recovery degree | 34.09 |
| | | Recovery rate | 27.27 |
| | | Recovery time | 15.91 |
| | | Multidimensional quantification | 9.09 |
| | | Latitude | 6.82 |
| | | Others | 6.82 |
| | Open herbaceous (n=93) | Recovery degree | 32.26 |
| | | Recovery rate | 22.58 |
| | | Others | 15.05 |
| | | Multidimensional quantification | 13.98 |
| | | Recovery time | 9.68 |
| | | Latitude | 6.45 |
| | Wetland (n=57) | Recovery rate | 38.60 |
| | | Recovery time | 21.05 |
| | | Recovery degree | 19.30 |
| | | Latitude | 10.53 |
| | | Others | 7.02 |
| | | Multidimensional quantification | 3.51 |

**Supplementary Table 7. Journals included in the literature synthesis to examine the usage of resilience in ecological research.** List of the 15 journals from which resilience-quantifying studies were identified. This list includes Nature, Science, Proceedings of the National Academy of Sciences due to their relevance in the field, as well as the 12 ecology journals with the highest Google Scholar h5-index at the time of data collection (January 2026).

| **Publication name** |
| --- |
| Nature |
| Science |
| Proceedings of the National Academy of Sciences |
| Nature Ecology & Evolution |
| Ecological Indicators |
| Ecology Letters |
| Molecular Ecology |
| Journal of Applied Ecology |
| Molecular Ecology Resources |
| Global Ecology and Biogeography |
| Journal of Ecology |
| Ecology |
| Ecography |
| Functional Ecology |
| Ecological Applications |

**Supplementary Table 8. Classification scheme for disturbance types.** Disturbance events reported in the literature were grouped into nine broad categories, each with associated subtypes (common examples shown). This classification was applied consistently during data extraction and subsequent analyses.

| Disturbance type | Subtypes (common examples) |
|---|---|
| Fire disturbance | Fire |
| Climatic disturbance | Drought; Temperature extremes; Precipitation extremes; Storm; Frost/Snow/Blizzard |
| Hydrological disturbance | Flood; Tsunami; Damming/Altered river flow |
| Geophysical disturbance | Landslide; Avalanche; Earthquake; Volcanic eruption; Soil erosion |
| Chemical disturbance | Nutrient loads (N, P, etc.); Heavy metal pollution; Salinity; Herbicide/Pesticide |
| Resource disturbance | Nutrient removal; Starvation |
| Biotic disturbance | Invasive species; Predator irruption; Disease; Herbivory (e.g., grazing); Bioturbation (e.g., burrowing) |
| Land-use and infrastructure development | Urbanisation; Road construction; Mining; Land conversion |
| Biological resource use | Logging; Plant harvesting; Fishing; Hunting |
| Structural disturbance | Population abundance manipulation; Population demographic manipulation; Community composition manipulation; Community interaction manipulation; Functional group manipulation |

**Supplementary Table 9. Overview of resilience-related frameworks and their key components.** Representative types of conceptual frameworks used in the reviewed studies, alongside their component names, common quantification approaches, and example references. Frameworks differ in how resilience is positioned conceptually: (1) frameworks in which resilience is decomposed into multiple components (*e.g.* recovery and resistance), (2) frameworks in which resilience and other components jointly form a broader stability concept, (3) frameworks in which resilience is presented alongside other measures without explicit integration, (4) "Ecosystem health" and "Ecological vulnerability" frameworks treat resilience as one of several components and do not include explicit recovery or resistance terms; these type of frameworks appeared only among landscape-level studies in our dataset. These frameworks were classified under a single "large-scale framework" category, though the table highlights the two most common types within this group.

<table>
<tr><th>Framework type</th><th>Component name</th><th>Quantify</th><th>Examples</th></tr>
<tr><td rowspan="3">Resilience: Recovery + Resistance + other components (optional)</td><td>Recovery</td><td>Degree of recovery / rate of recovery / recovery time</td><td rowspan="3">Mulla et al., 2024[1]; Perez et al., 2024[2]</td></tr>
<tr><td>Resistance</td><td>Resistance</td></tr>
<tr><td colspan="2">Optional components: invariability, resilience, relative resilience, compensation, etc.</td></tr>
<tr><td rowspan="3">Stability: Resilience + Resistance + other components (optional)</td><td>Resilience</td><td>Degree of recovery / rate of recovery / recovery time</td><td rowspan="3">Ding et al., 2023[3]; Zhang & Want, 2023[4]</td></tr>
<tr><td>Resistance</td><td>Resistance</td></tr>
<tr><td colspan="2">Optional components: recovery, invariability</td></tr>
<tr><td rowspan="3">Unintegrated: Resilience + Resistance + Recovery (optional)</td><td>Resilience</td><td>Degree of recovery / rate of recovery</td><td rowspan="3">Kokkonen et al., 2024[5]; Wallace & Walsworth, 2024[6]</td></tr>
<tr><td>Resistance</td><td>Resistance</td></tr>
<tr><td>Recovery</td><td>Degree of recovery / rate of recovery</td></tr>
<tr><td rowspan="4">Ecosystem health: Vigor + Organisation + Resilience + other components (optional)</td><td>Vigor</td><td rowspan="2"></td><td rowspan="4">Lin et al., 2024[7]; Shengrui et al., 2024[8]</td></tr>
<tr><td>Organisation</td></tr>
<tr><td>Resilience</td><td>Inferred by environmental context</td></tr>
<tr><td colspan="2">Optional components: service, pressure, contribution.</td></tr>
<tr><td rowspan="3">Ecological Vulnerability: Sensitivity + Resilience + other components (optional)</td><td>Sensitivity</td><td></td><td rowspan="3">Chen et al., 2024[9]; Shi et al., 2023[10]</td></tr>
<tr><td>Resilience</td><td>Inferred by environmental context / rate of recovery</td></tr>
<tr><td colspan="2">Optional components: pressure</td></tr>
</table>

### S2 Supplementary Figures

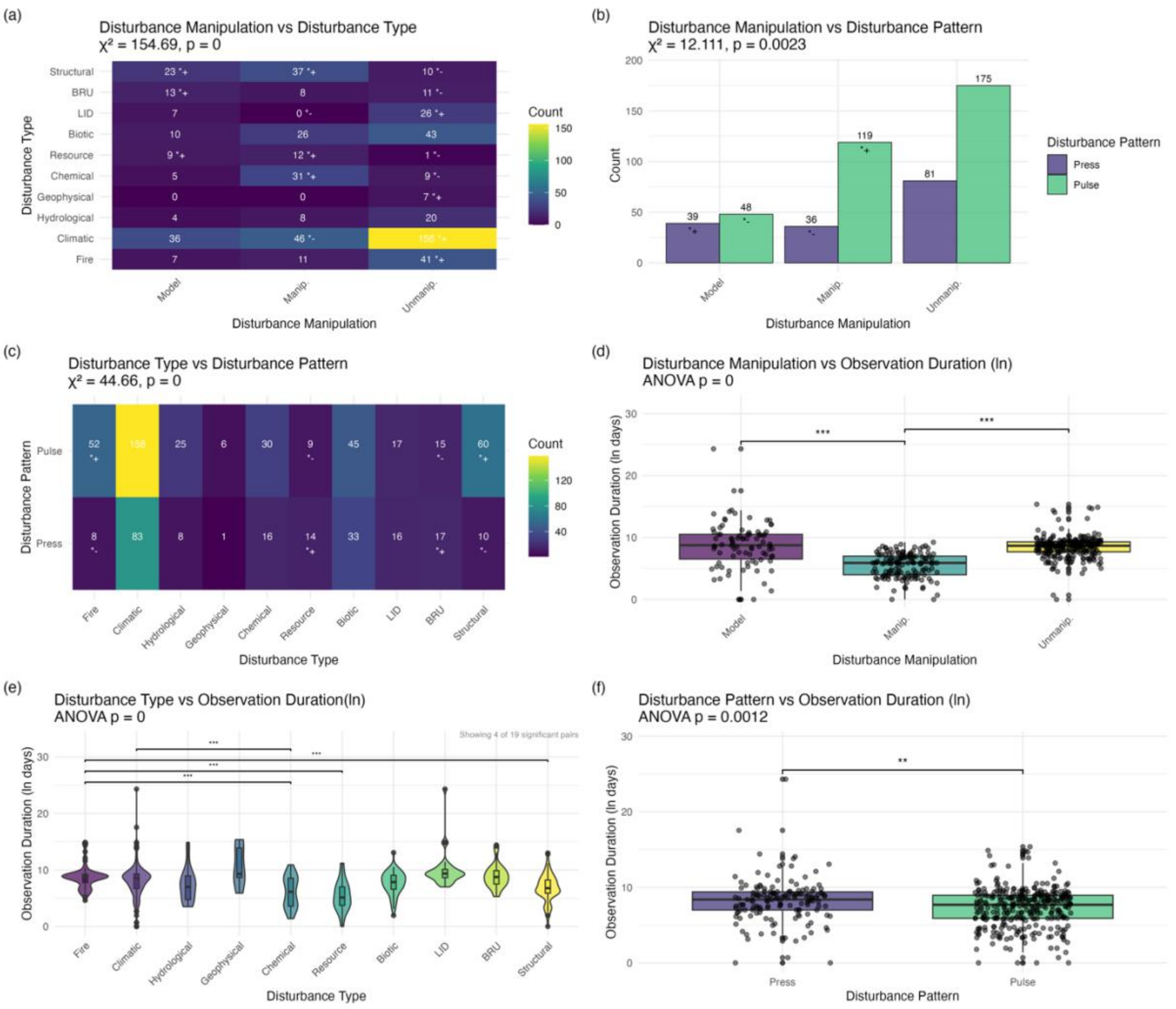


**Supplementary Fig. 1. Associations among disturbance-related study attributes.** Panels (a–c) show contingency analyses among disturbance manipulation, disturbance type and disturbance pattern across the analysed studies. Tile colour intensity and bar height represent study counts. Numbers within tiles or bars indicate counts; symbols denote significant deviations from independence based on standardised residuals from chi-squared tests (|residual| > 1.96). (a) Disturbance manipulation versus disturbance type. (b) Disturbance manipulation versus disturbance pattern. (c) Disturbance type versus disturbance pattern. Panels (d–f) compare log10-transformed observation duration (days) across disturbance categories using one-way ANOVA followed by Tukey's HSD post-hoc tests where appropriate. Points represent individual studies. (d) Observation duration across manipulation categories. (e) Observation duration across disturbance types; for clarity, only the four most significant pairwise contrasts are displayed. (f) Observation duration between disturbance patterns (Press and Pulse). Brackets indicate significant pairwise contrasts; asterisks denote adjusted P values (* P < 0.05; ** P < 0.01; *** P < 0.001; **** P < 0.0001).

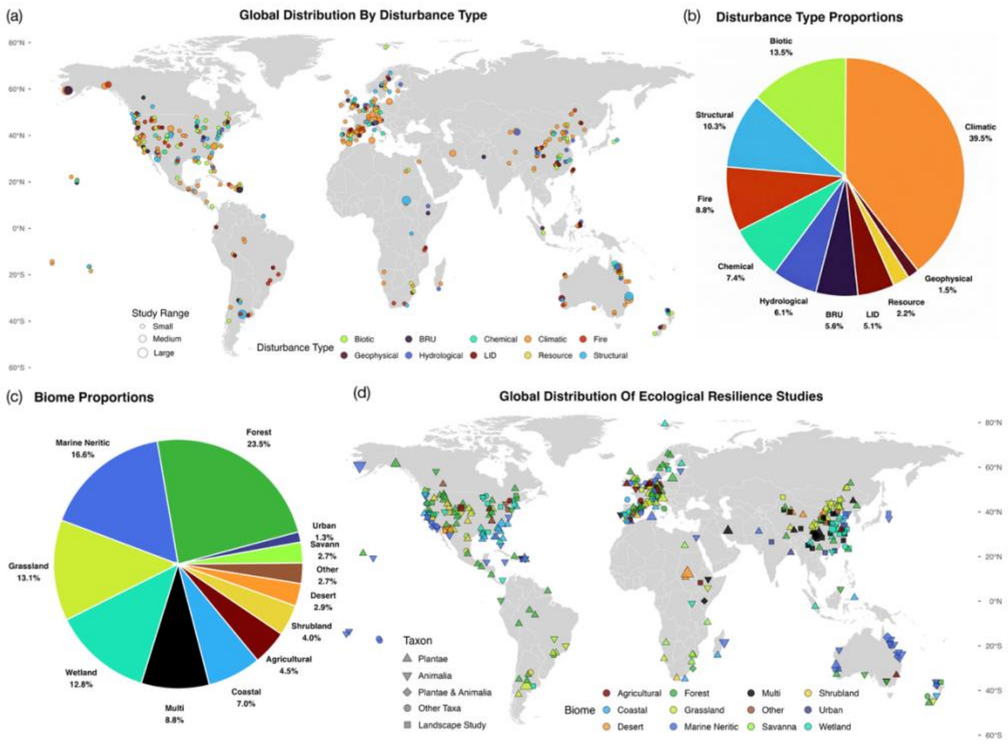


**Supplementary Fig. 2. Global distribution of resilience studies by disturbance type, biome, and focal taxa, together with their proportional representation.** (a) Global distribution of field studies by disturbance type. Each point marks a study location, coloured by disturbance type. Point size indicates the study range. (b) Proportion of studies including each disturbance type. (c) Global distribution of field studies by biome and focal taxon. Points are coded by biome (colour) and focal taxon (triangle = plants, inverted triangle = animals, diamond = including both plants and animals, circle = other taxa, square = landscape studies). Point size indicates the study range. (d) Proportions of studies by biome. The map uses an equal-aspect view and excludes Antarctica (displayed latitude −60° to 80°). In this figure, biome corresponds to the habitat type categories used throughout the study.

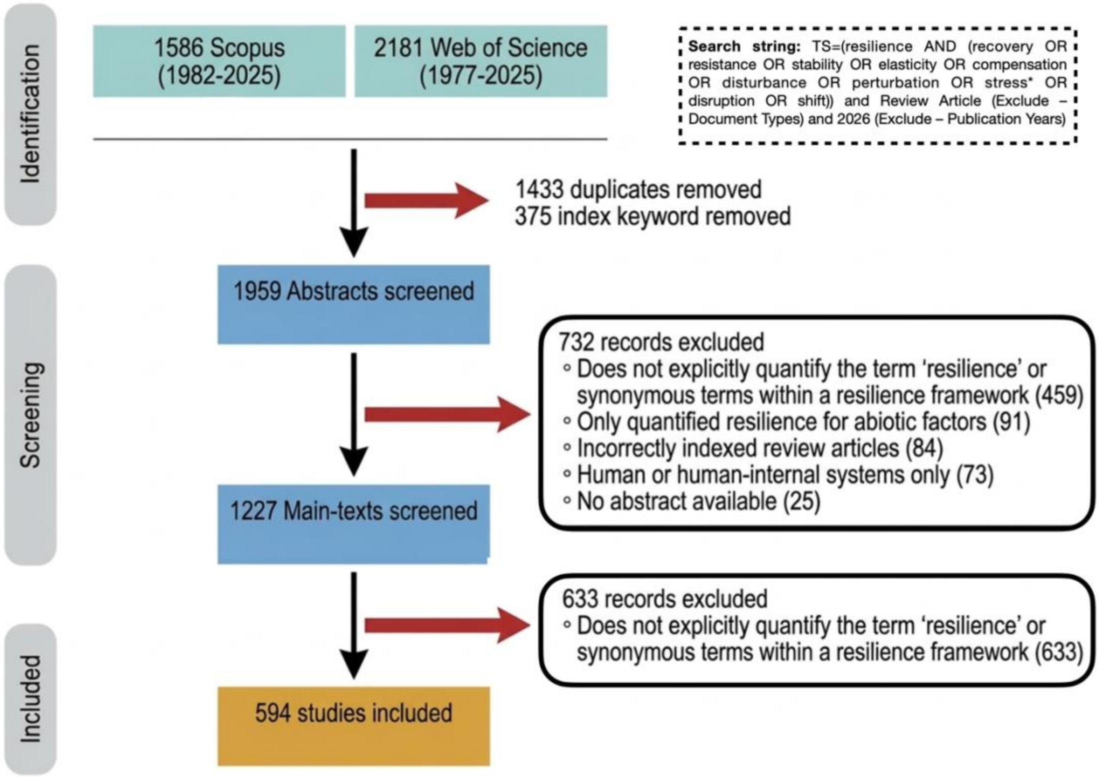


**Supplementary Fig. 3. PRISMA diagram describing the search results in different search engines and the different steps of selecting articles for inclusion in the systematic review.** Depicted are the number of studies excluded at each stage. "Index keywords removed" refers to excluding articles in which the specified keywords appear only in indexing keywords, but not in the title, abstract, or author-provided keywords. In this figure, 'main-texts screened' corresponds to the 'full-text screened' used throughout the study.

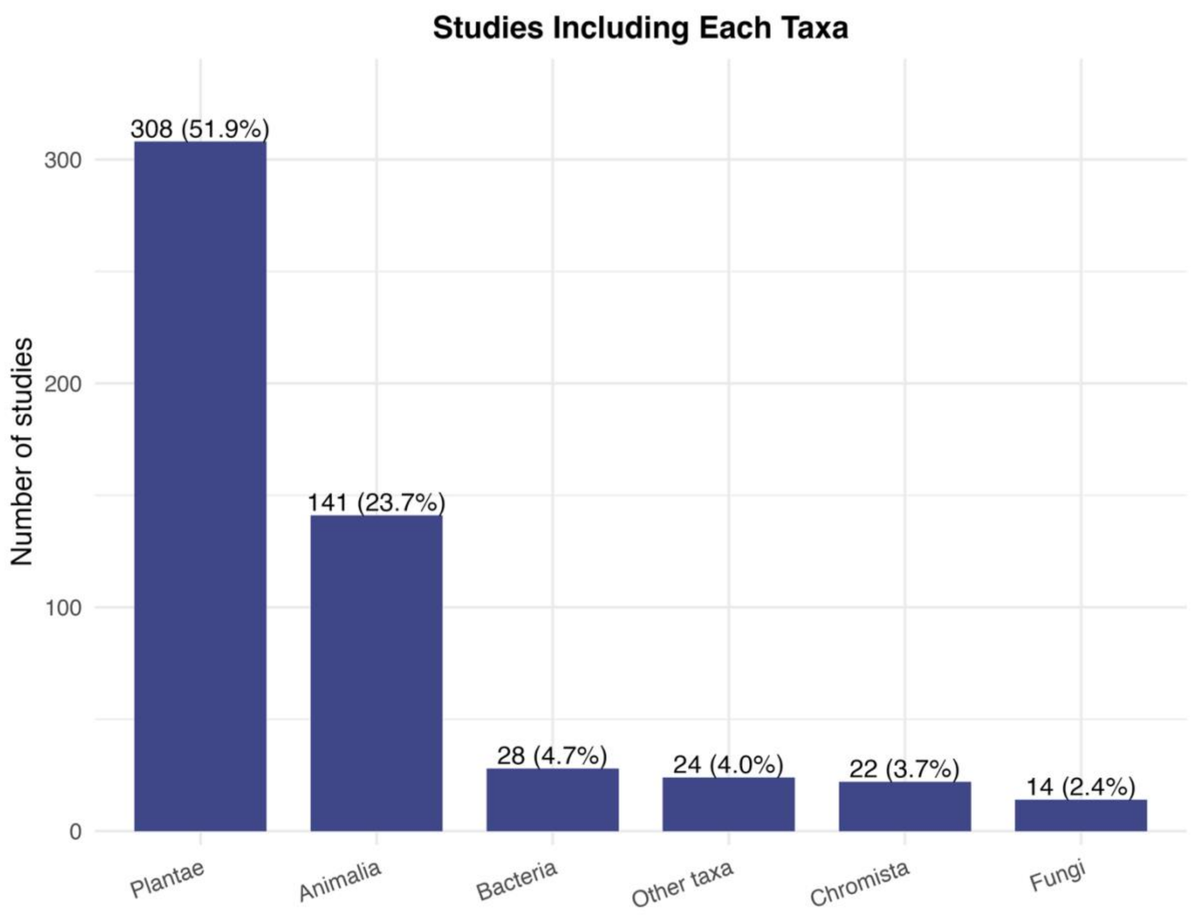


**Supplementary Fig. 4. More than half of the studies examined subjects from Plantae.** Number of studies including different taxonomic groups. “Other Taxa” includes groups that span multiple kingdoms, such as phytoplankton, benthic organisms, and virtual species, which refers to model-based studies using hypothetical species without clear taxonomic affiliation. Numbers above bars show study counts with percentages in brackets. Percentages are calculated relative to the total number of studies, so their sum exceeds 100% because many studies include multiple taxa.

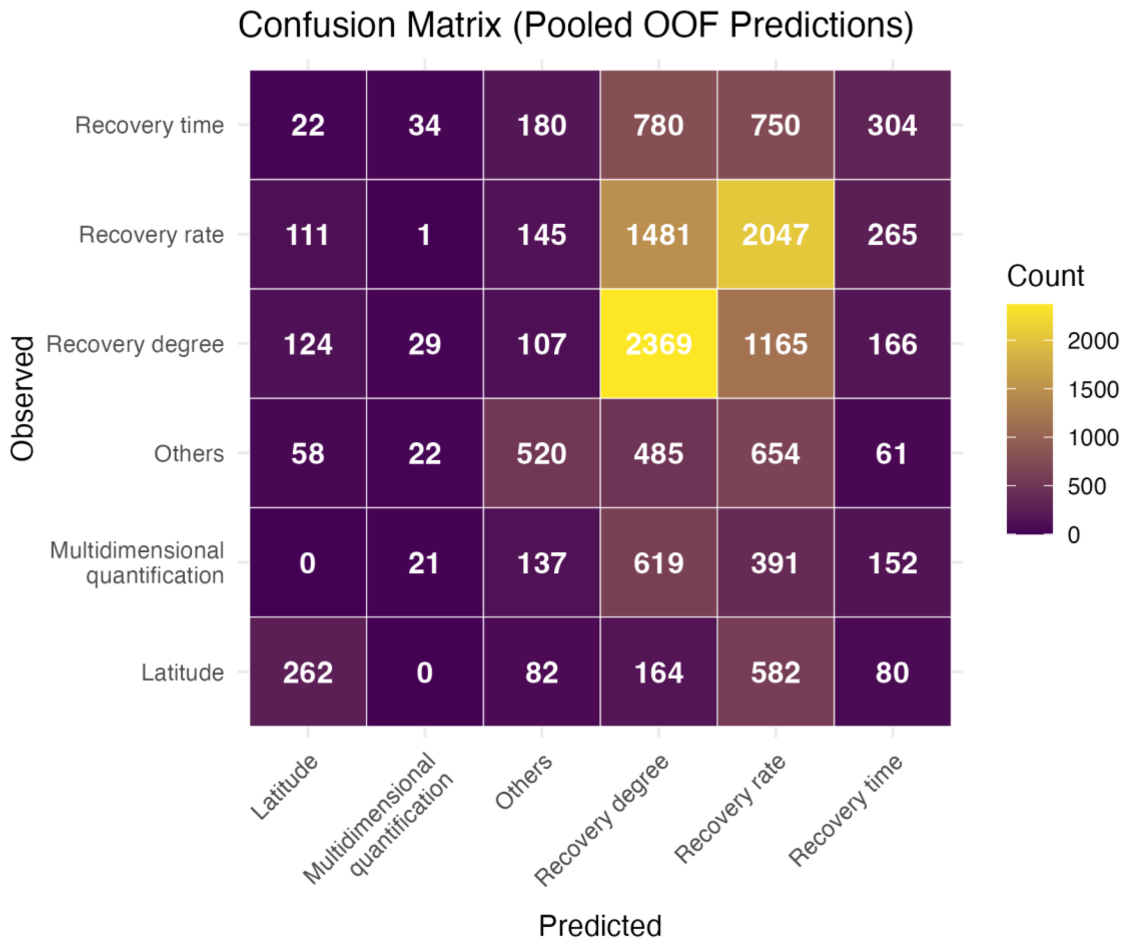


**Supplementary Fig. 5. Recovery time is often confused with recovery degree and recovery rate.** Confusion matrix of resilience-metric classification from cross-validated model predictions. The heatmap shows the confusion matrix derived from pooled out-of-fold (OOF) predictions of the conditional inference forest used to classify resilience-quantification categories. Rows represent the observed resilience metric reported in each study, and columns represent the metric predicted by the model. Cell values indicate the number of predictions aggregated across all cross-validation runs. Correct classifications appear along the diagonal, whereas off-diagonal cells indicate misclassification among metric categories.

### S3 Supplementary Methods

**Details of data collection**

To examine how resilience is quantified in ecology and how study context shapes metric choice, we extracted a standard set of study attribute variables from the 594 articles that met our inclusion criteria (see Supplementary Data 1). We extracted *Resilience quantification* (see Table 1), *i.e.*, how resilience is quantified, as the objective of this study is to characterise how resilience has been operationalised in empirical ecological research. We also extracted other study attribute, including (1) the *measure variable type* for resilience quantification (see Extended Data Table 1), since the choice of what is measured is central to study design[11]; (2) the *level of biological organisation* (*i.e.*, individual, population, community, ecosystem, and landscape), to test for the potential to scale resilience (See Discussion) and the how different fields may quantify resilience differently; (3) the *approach* applied (*i.e.*, modelling, laboratory experiment, field experiment, and field observation), to account for evaluate how experimental control affects the quantification of resilience; (4) the *disturbance type* (*e.g.*, drought, fire; see Extended Data Table 2), to assess whether certain disturbance types favour particular resilience metrics; (5) the *disturbance pattern* (categorised as pulse [short-term discrete events] *vs.* press [long-term continuous pressures][12]), because mechanism-specific disturbances plausibly favour different metrics; (6) the *observation duration* (the interval between the first and last observations), because the time window of the study may condition the detectability of different resilience components; (7) the focused *habitat type*[13], because different ecosystems may favour different resilience metrics; (8) The focused *taxonomic group*, because inherent differences among taxa may imply that the most appropriate resilience metrics vary across systems[14,15]; (9) the *resilience framework* employed (*e.g.*, stability, ecosystem health; see Supplementary Table 2), defined as how studies operationalise resilience either by using multiple metrics to represent it, or by treating resilience as a single metric embedded within a broader conceptual framework; (10) the *study location*, to help represent the geographical distribution of studies; (11) the *approach to apply disturbances* (*i.e.*, model, manipulated, unmanipulated), to help represent the degree of experimental control over disturbances in the studies; (12) the *country of the first author's institution,* to examine whether geographical academic networks might influence the choice of metric in the explore analysis.

To facilitate the collection of these data, we classified the studies into direct-response *vs.* inference-based studies. We based this distinction on how resilience was measured: direct-response studies quantify resilience from measurements taken directly on the system's response to disturbance[16]. Inference-based studies infer resilience from indicators or models that act as proxies rather than direct response measurements[8].We coded disturbance-related variables (*e.g.*, disturbance type, observation duration) for direct-response studies only, as such information was typically unavailable or ambiguous in inference-based studies. To enable consistent comparisons across resilience studies, we first standardised the diverse terminology used to quantify resilience in the literature. Reported metrics were harmonised into six quantitative categories—recovery rate[17,18], recovery degree[19–21], recovery time[22,23], resistance[24–26], latitude[27–29] and invariability[2,30,31]—each assigned to a single term (see Table 1; The citations here are the representative examples of each quantitative categories). Metrics of resilience that did not align with these categories are classified as "other". Importantly, we note that a given study may employ multiple metrics.

**Details of LLM-related task**

To assist with literature screening and structured data extraction, we implemented a large language model (LLM) workflow comprising three tasks: abstract screening, full-text screening, and study-attribute extraction. All prompts used in these tasks and the performance of successive prompt versions during development and validation are provided in Supplementary Data 1.

Following the literature search, we conducted LLM-assisted screening in two stages (Supplementary Fig. 1). First, the abstracts of the retrieved articles were screened to identify studies likely to quantify ecological resilience. Abstract text was provided to the model in CSV format, and the model evaluated predefined inclusion criteria through structured questions. Responses were returned as compact JSON outputs and parsed into tabular format.

Articles passing abstract screening proceeded to full-text screening. At this stage, article PDFs were parsed using the PyPDF2 package and submitted individually to the model through the OpenAI API. The model evaluated the same inclusion criteria using the full article text. Articles satisfying both stages were retained for data extraction. We provide the complete lists of initially retrieved articles, abstract-screened articles, and the final 594 included studies in Supplementary Data 2.

We developed prompts for each task iteratively using human-coded reference datasets. Initial prompt versions were tested on training subsets of manually coded articles. We then examined model outputs to identify systematic errors and revised the prompts accordingly. For example, early prompt versions frequently classified studies using remote-sensing data as inference-based even when these data represented direct measurements of ecological variables. Instructions were therefore refined to clarify the distinction between direct-response and inference-based measurements.

We evaluated prompt performance against human annotations. For screening tasks, we quantified agreement using Cohen's κ and overall consistency. For abstract screening, we additionally calculated the false-negative rate relative to human decisions. These metrics were computed by comparing LLM outputs with manually coded reference data. For the data-extraction task, model performance was assessed using F1-scores comparing extracted attributes with human-coded values across study variables. These scores were calculated after standardising categorical outputs and applying post-processing rules to harmonise terminology across model outputs and human annotations. Finally, we conducted validation on independent article subsets separate from those used for prompt development. Only prompts meeting the predefined performance thresholds described in Methods were deployed to the remaining articles.

After each deployment stage, we conducted an additional audit to verify that model performance remained consistent when applied to previously unseen articles. For each task, a random sample of 50 articles that had not been used during prompt development or validation was manually checked and compared with the model outputs. Agreement metrics were recalculated to confirm that performance remained above the predefined thresholds. This procedure was implemented after abstract screening, full-text screening, and data extraction. The resulting performance metrics from all prompt versions and auditing steps are summarised in Supplementary Data 1.

For the final set of included studies, the LLM was instructed to extract predefined study attributes as structured JSON outputs. Each prompt contained a series of questions corresponding to individual variables, including resilience quantification, measured variable type, disturbance characteristics, and study context. Parsed outputs were converted into tabular format and subjected to automated post-processing to standardise categorical labels and resolve formatting inconsistencies. Because ecological studies frequently report multiple values for one study attribute variables, such as resilience metrics, extraction variables allowed multiple values separated by delimiters. Post-processing procedures subsequently harmonised these values and mapped them to the predefined classification scheme used in our analyses.

Although all extraction variables met the predefined F1-score thresholds during validation, we detected a systematic bias in the extraction of resilience quantification. Specifically, when studies adopted multidimensional resilience frameworks (see Supplementary Table 2), the model often returned only a single quantification metric even when multiple metrics were explicitly reported. To address this issue, we manually reviewed all direct-response studies classified as belonging to a resilience framework but assigned a single quantification value by the model (n = 65). This review resulted in the addition of previously omitted quantification metrics for 25 studies. Because resilience quantification constitutes the primary response variable in our analyses, we produced two versions of the final dataset. The first dataset consists of the direct outputs of the LLM following automated post-processing. The second dataset incorporates the manual corrections described above, which affect only the resilience-quantification variable while leaving all other attributes unchanged. Providing both datasets ensures transparency while minimising potential bias introduced by manual intervention. The LLM-only dataset preserves the fully automated extraction pipeline, whereas the corrected dataset resolves the identified systematic omission in resilience quantification. Both datasets, together with the manually curated subsets used during prompt development and validation, are provided in Supplementary Data 3.

**Details of conditional inference forest**

To test whether study context predicts the resilience quantification used by authors, we fitted conditional inference forests (CIFs)[32,33] to studies that directly quantified resilience from observed system responses. The response variable was quantification category, and the main model included eight predictors: disturbance pattern, measured variable, approach, ecological level, disturbance type, taxon, habitat type, and observation duration. Observation duration was treated as a continuous predictor after $\log_{10}$-transformation; all other predictors were categorical. The modelling workflow and data-processing logic followed the analysis scripts provided for this study.

Because several predictors contained sparse categories, missing values, or multi-label annotations, predictors were harmonised before model fitting. Small categories were merged when they were conceptually adjacent and, where relevant, showed similar distributions across quantification categories. Missing values were manually checked where possible and otherwise imputed conservatively when they reflected conceptual non-comparability rather than unknown true values, as in a small subset of studies lacking a real-world observation duration. For predictors with multi-label annotations, we retained a single category using explicit priority rules designed to preserve the aspect judged most likely to constrain or dominate the resilience quantification used. For example, for approach, we defined the priority ranking modelling-based simulation > field experiment > field observation, reflecting decreasing levels of experimental control. When a study combined modelling and field observation, we therefore retained modelling-based simulation, as the higher level of control was assumed to impose stronger constraints on the form of quantification applied. These priority rules were applied to measured variable, approach, ecological level, disturbance type, and disturbance pattern. For disturbance type, structural labels were removed when they co-occurred with one or more concrete disturbance types, because they describe an abstract level of manipulation rather than a real-world disturbance class. Full details of category consolidation, manual corrections, and priority rules are provided in the analysis scripts.

Hyperparameters were tuned using repeated stratified cross-validation with macro-F1 as the primary performance metric. We first assessed sensitivity to mincriterion, the split threshold in the conditional inference framework, and compared values of 0, 0.9, and 0.95. We used mincriterion = 0 in the main analysis because it gave the highest predictive performance, and retained mincriterion = 0.9 for robustness testing. We next tuned mtry from 1 to 6 and selected mtry = 3 using the 1-SE rule[34], which was also consistent with the square root of $p$ heuristic for classification forests[35]. We then evaluated ntree from 250 to 4,000 and selected 2,000 trees for the main model because performance had stabilised across this range and peaked at this value. Permutation-based variable importance was estimated using 50 permutations per predictor per cross-validation fold. All tuning results are reported in Supplementary Data 4.

We also fitted alternative CIFs to assess the sensitivity of the analysis to data-processing and model-specification choices. These comprised a separate set of nine robustness models and an exploratory model that additionally included journal and institution country. The robustness set included seven alternative encodings in which priority rankings used to collapse multi-label predictors were reversed, one model excluding the multidimensional quantification response class, and one model using *mincriterion* = 0.9 instead of 0. The results of these alternative CIFs are reported in the Supplementary Results.

### S4 Supplementary Results

#### Sensitivity tests of conditional inference forest

We fitted nine alternative conditional inference forests (CIFs) to test the sensitivity of the main analysis to predictor harmonisation rules and model specification. Across all nine runs, the overall pattern of variable importance was highly consistent (Supplementary Fig. 4). Measured variable, approach and disturbance pattern were always the three most important predictors, whereas taxon, habitat type, and usually disturbance type contributed little to predictive performance.

Reversing the priority rankings used to collapse multi-label predictors caused only modest changes in mean macro-F1 and did not alter the substantive interpretation of the model. The most pronounced change among these rule-based alternatives occurred when both pulse- and press-disturbance rankings were reversed, although even here the dominant importance of measured variable, approach and disturbance pattern was retained. Removing the multidimensional quantification class substantially increased mean macro-F1, as expected from the simplified classification task after exclusion of one, comparatively heterogeneous, response category. However, the overall ranking of predictor importance remained unchanged, indicating that the main CIF conclusions were robust to the inclusion of this class. Moreover, increasing the split threshold to *mincriterion* = 0.9 had little effect on variable-importance ranking and only a small effect on performance. Together, these results indicate that the main CIF conclusions are robust to alternative encoding rules and parameter settings, and that part of the predictive difficulty of the full model reflects heterogeneity in the response variable itself.

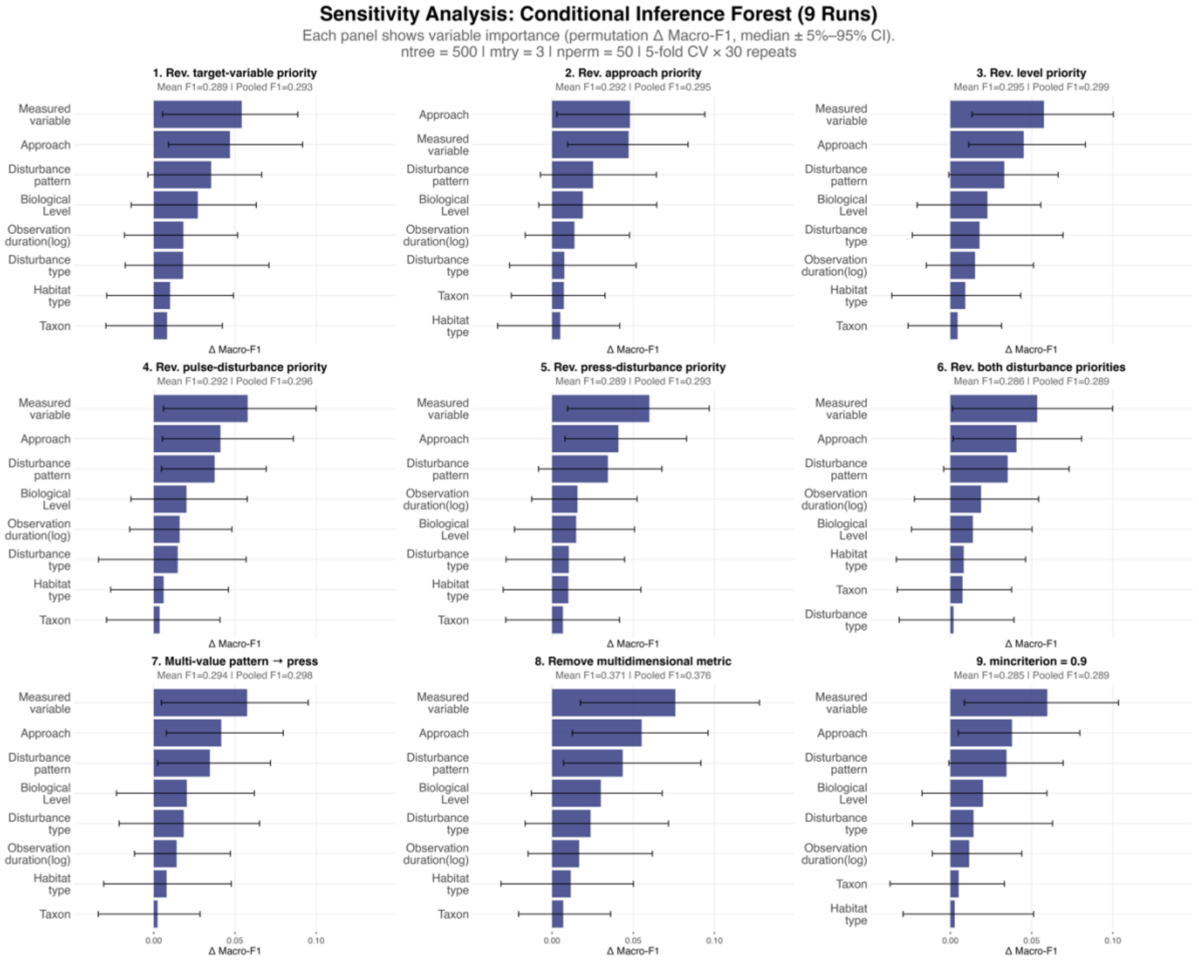


**Supplementary Fig. 6. Nine alternative conditional inference forests identified the same top three predictors as the main model, supporting the robustness of the main results.** Variable-importance profiles for nine alternative conditional inference forests (CIFs) used to test the robustness of the main analysis. The nine runs comprised reversed priority rules for measured variable, approach, biological level, pulse-disturbance type, press-disturbance type, both disturbance-type rankings, and multi-value disturbance pattern assignment, together with one model excluding the multidimensional quantification response class and one model using a stricter split threshold (mincriterion = 0.9). In each panel, bars show median permutation importance for each predictor, expressed as the decrease in macro-F1 after permutation, and error bars indicate the 5th–95th percentile interval across repeated cross-validation runs. Panel subtitles report mean cross-validated macro-F1 and pooled out-of-fold macro-F1. In this figure, 'measured variable' corresponds to the predictor termed 'target variable group' elsewhere in the study.

**An alternative conditional inference forest for exploration**

To explore whether publication venue and institutional geography added predictive structure beyond the main contextual predictors, we fitted an alternative CIF. In addition to the variables used in the main model, this exploratory analysis included journal and institution country. We excluded these variables from the main analysis because they do not directly describe study design or ecological context, but they may still capture broader disciplinary or institutional patterning in how authors quantify resilience. As in the main analysis, we harmonised institution country into broad regional groups and merged journals with small sample sizes into broader scope-based categories before modelling.

Adding these predictors did not change the overall interpretation of the model (Supplementary Fig. 5). Measured variable (*i.e.,* target variable in the figure) remained the strongest predictor, followed by approach and disturbance pattern. Journal added a moderate amount of predictive signal and ranked fourth, whereas institution continent contributed little and ranked below disturbance type, ecological level, and observation duration. Habitat type and taxon again remained the weakest predictors. In repeated resampling runs, target variable ranked among the top three predictors in 88% of runs and had positive permutation importance in 99% of runs; the corresponding values were 65% and 94% for approach, 48% and 93% for disturbance pattern, and 46% and 83% for journal. By contrast, institution continent ranked among the top three predictors in only 2% of runs and showed positive importance in 68% of runs. These results suggest that publication venue captures some additional variation in quantification practice, whereas institutional geography at this broad scale adds little.

Overall, the exploratory CIF supports rather than revises the main conclusion. Study context still explains only a limited share of the variation in resilience quantification, and journal-level differences appear secondary to the main contextual predictors identified in the primary analysis.

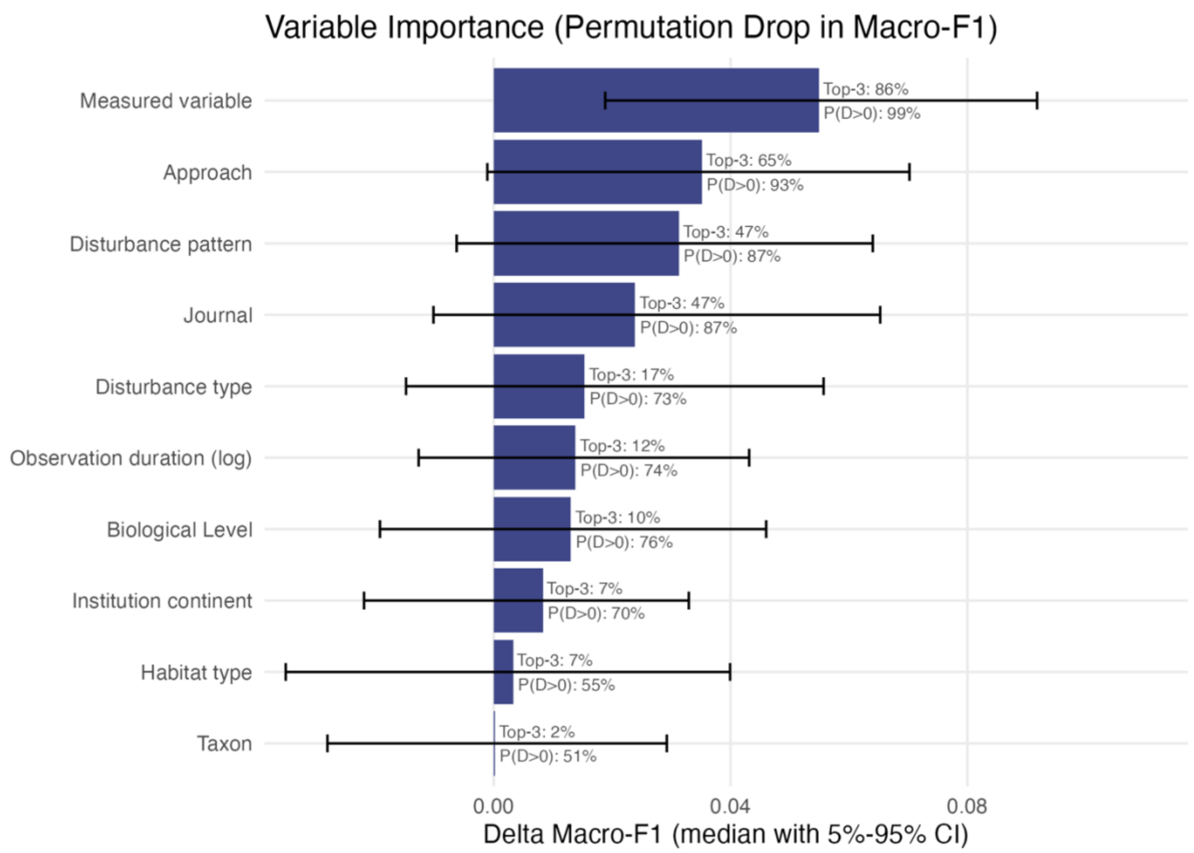


**Supplementary Fig. 7. An exploratory conditional inference forest including journal and institution continent identified the same top three predictors as the main model.** Variable-importance profile from an exploratory conditional inference forest (CIF) used to assess whether publication venue and institutional geography add predictive structure beyond the main contextual predictors. In addition to the predictors included in the main model, this analysis incorporated journal and institution country, with the latter harmonised into broad continental groups and shown here as Institution continent. Bars show median permutation importance for each predictor, expressed as the decrease in macro-F1 (ΔMacro-F1) after permutation, and horizontal error bars indicate the 5th–95th percentile interval across repeated cross-validation runs. Text annotations report the proportion of runs in which each predictor ranked among the three most important predictors (Top-3) and the proportion of runs in which permutation importance was positive (P(Δ>0)). The exploratory model retained the same dominant predictors as the main analysis, with target variable, approach, and disturbance pattern contributing the most predictive signal. Journal added moderate importance, whereas institution continent, habitat type, and taxon contributed little. In this figure, ‘Target variable’ corresponds to the predictor termed ‘measured variable’ elsewhere in the study.

**Results from half of the journals**

To evaluate the robustness of our conclusions to journal selection, we assessed whether the main conclusions of this study depend on the set of journals included by repeating key analyses using two subsets of the data: the first half of the selected journals and the remaining journals. Because the conditional inference forest requires a larger sample size to achieve stable performance, we restricted this comparison to the analyses corresponding to the first three main-text figures.

Across these comparisons, the most pronounced difference between the two subsets was the prevalence of inference-based studies. In the first-half journals, inference-based studies accounted for 37.6% of the dataset, compared with only 7.7% in the remaining journals (Supplementary Figs. 6 and 7). Consistent with this difference, the most common resilience quantification also differed between the two subsets: recovery rate dominated in the first-half journals, whereas recovery degree was most frequent in the remaining journals. These results indicate that conclusions about which metric is most commonly used are sensitive to journal selection and may be biased when based on a limited subset of journals.

Despite this difference in ranking between recovery rate and recovery degree, the broader patterns remained consistent with the main analysis. In both subsets, resilience quantification showed no clear convergence over time, remained concentrated within a small number of commonly used categories, and exhibited an increasing use of multidimensional quantification in recent years (Supplementary Fig. 7). Thus, apart from the relative ordering of the two most common metrics, the overall structure of quantification practices was robust to journal subsampling.

The Sankey diagrams further show that the relationships among ecological level, approach, and measured variable remained similarly complex in both subsets (Supplementary Fig. 8). However, the first-half journals included a greater proportion of studies that measured environmental context variables at the landscape level. Because most of these studies were inference-based (92.5%), this difference largely explains the higher proportion of inference-based studies observed in the first-half journals and, consequently, the differences seen in Supplementary Figs. 6 and 7. Moreover, the distribution of research scope across journals provides further support for this interpretation (Supplementary Fig. 9). Most landscape-level environmental context studies originated from *Ecological Indicators* (n = 40), with a smaller contribution from *Proceedings of the National Academy of Sciences of the United States of America* (n = 6). This concentration indicates that journal scope can strongly shape the types of resilience metrics reported, thereby influencing aggregate patterns when only a subset of journals is considered.

Taken together, these results show that journal-specific differences, particularly in scope, can affect the apparent prevalence of individual resilience metrics. However, the main conclusions of the study remain unchanged: resilience quantification does not converge over time, is dominated by a small set of commonly used metric categories, and has diverse and cross-level study context.

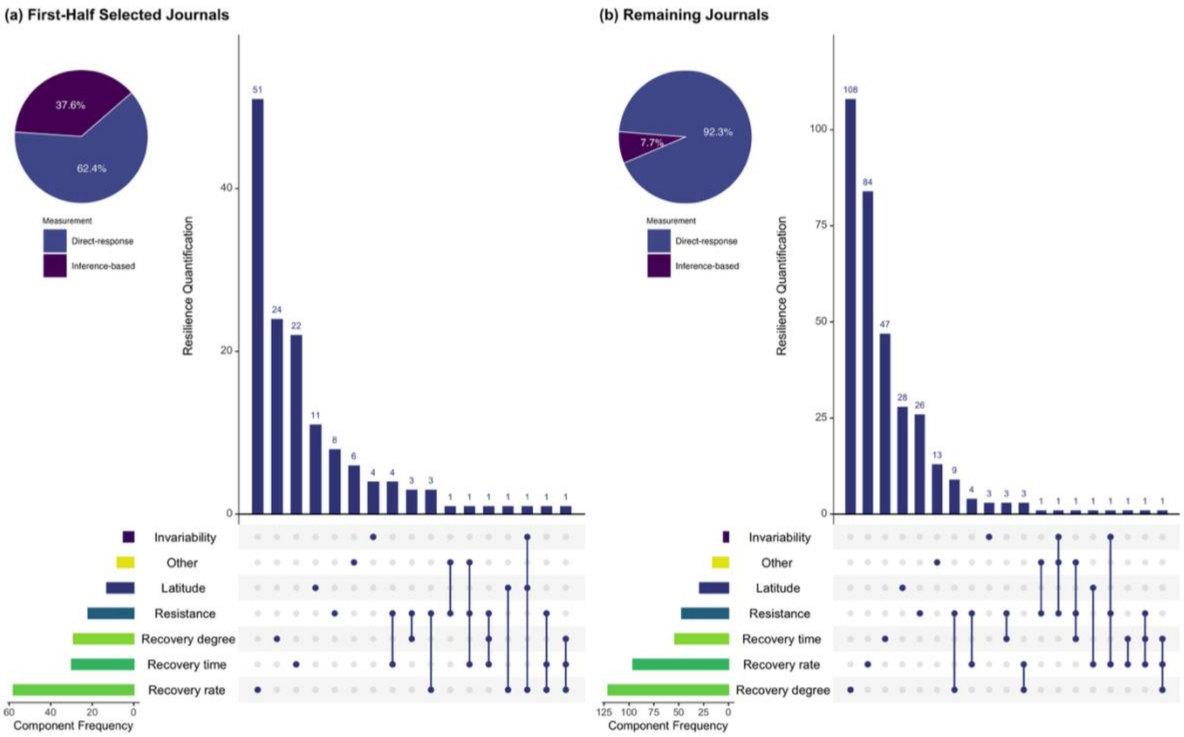


**Supplementary Fig. 8. Commonly used resilience metrics are ranked differently between the first-half selected journals and the remaining journals.** The pie chart shows the proportion of direct-response and inference-based studies. The UpSet plot shows the frequency of individual quantification components and their combinations; bars represent the number of studies using each combination, and connected dots indicate co-occurring components within the same study. (b) Same as (a), but for the remaining journals.

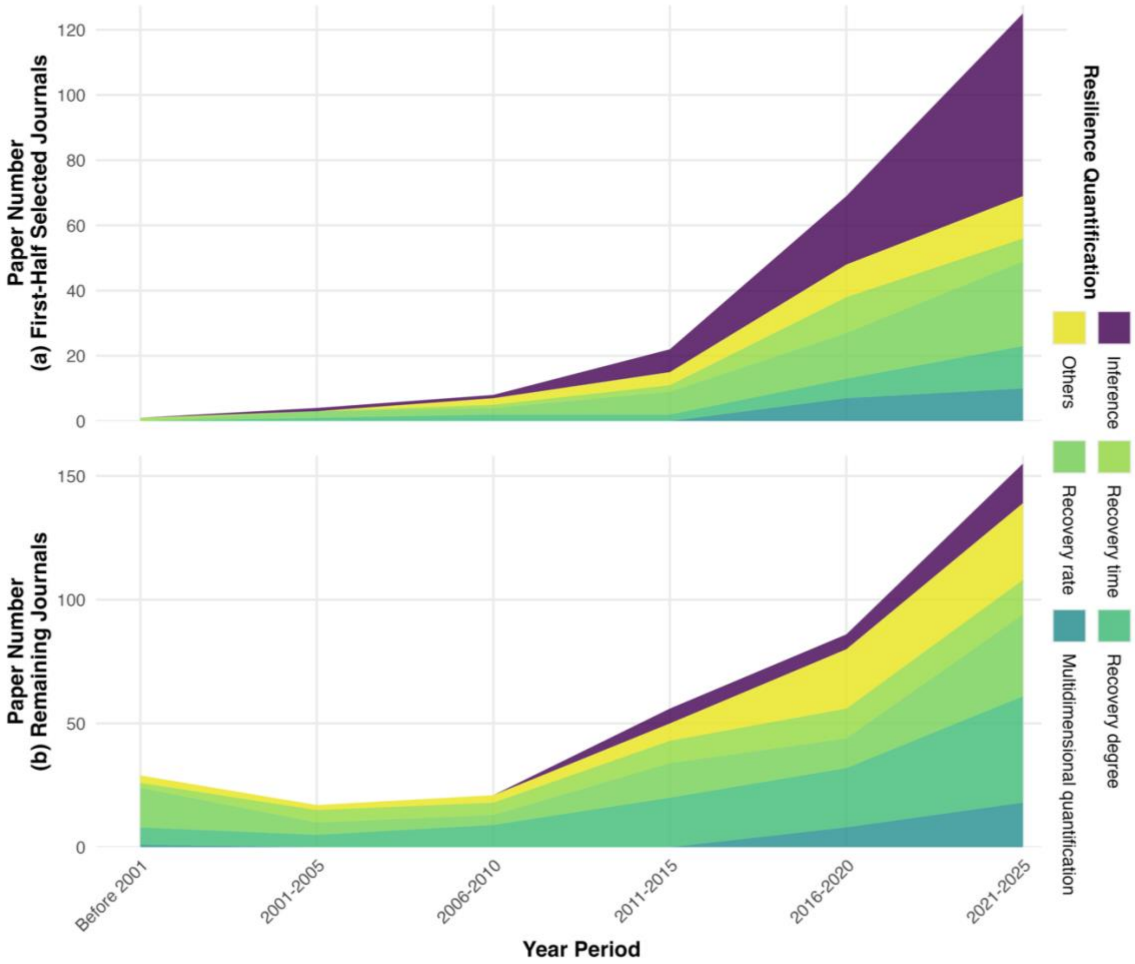


**Supplementary Fig. 9. Temporal trends in resilience quantification show no convergence across either journal subset, with the primary difference being a higher prevalence of inference-based studies in the first half.** (a) Temporal trends in the number of studies using different resilience quantification categories in the first-half selected journals. (b) Same as (a), but for the remaining journals. Shaded areas represent the number of studies within each quantification category across time periods. In both subsets, resilience quantification remains concentrated within a limited set of categories and shows no clear convergence over time.

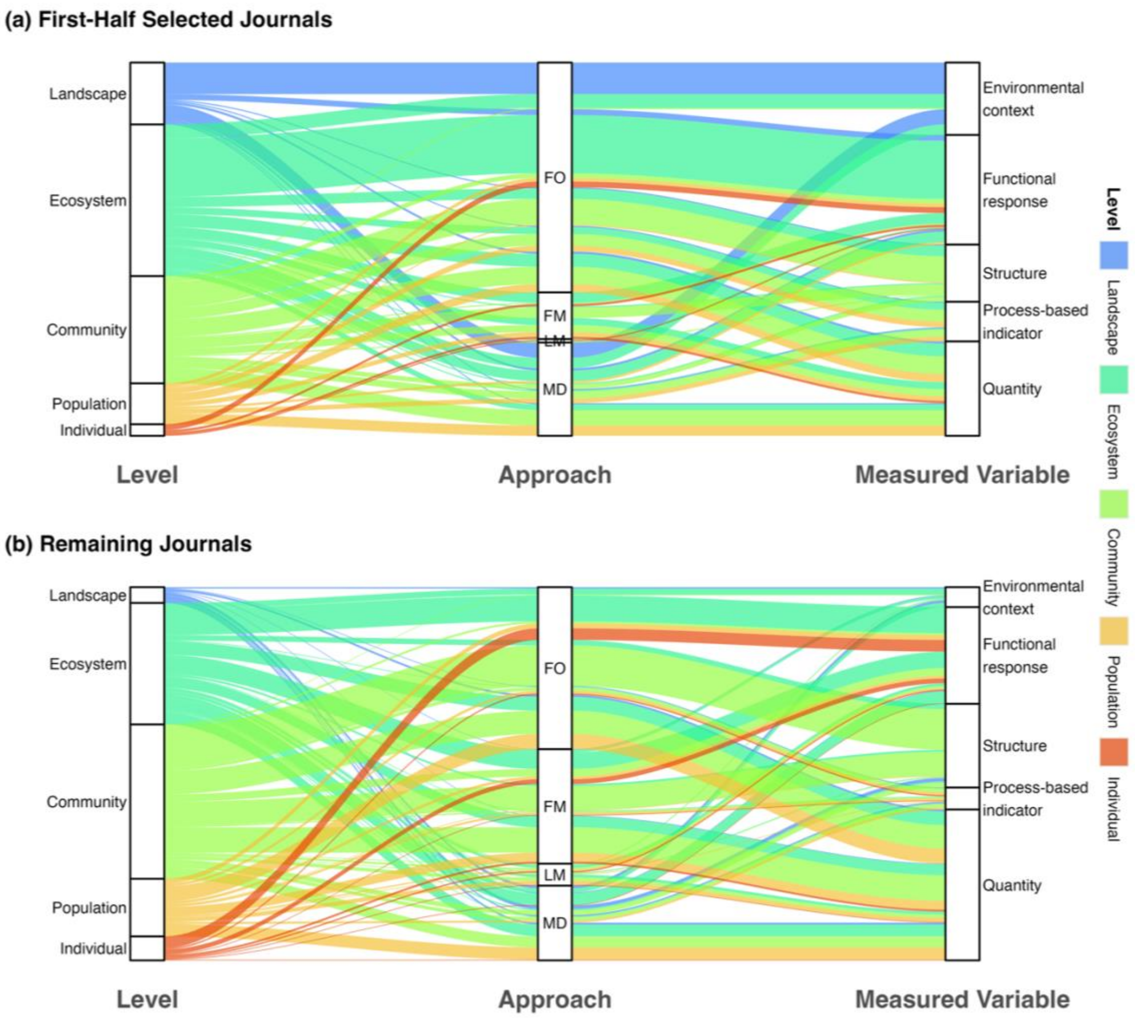


**Supplementary Fig. 10. Relationships among ecological level, approach, and measured variables are diverse across both journal subsets, but landscape-level studies are more prevalent in the first half.** (a) Alluvial diagram showing the relationships among ecological level, approach, and measured variable for the first-half selected journals. Flows represent the number of studies linking categories across the three dimensions. (b) Same as (a), but for the remaining journals.

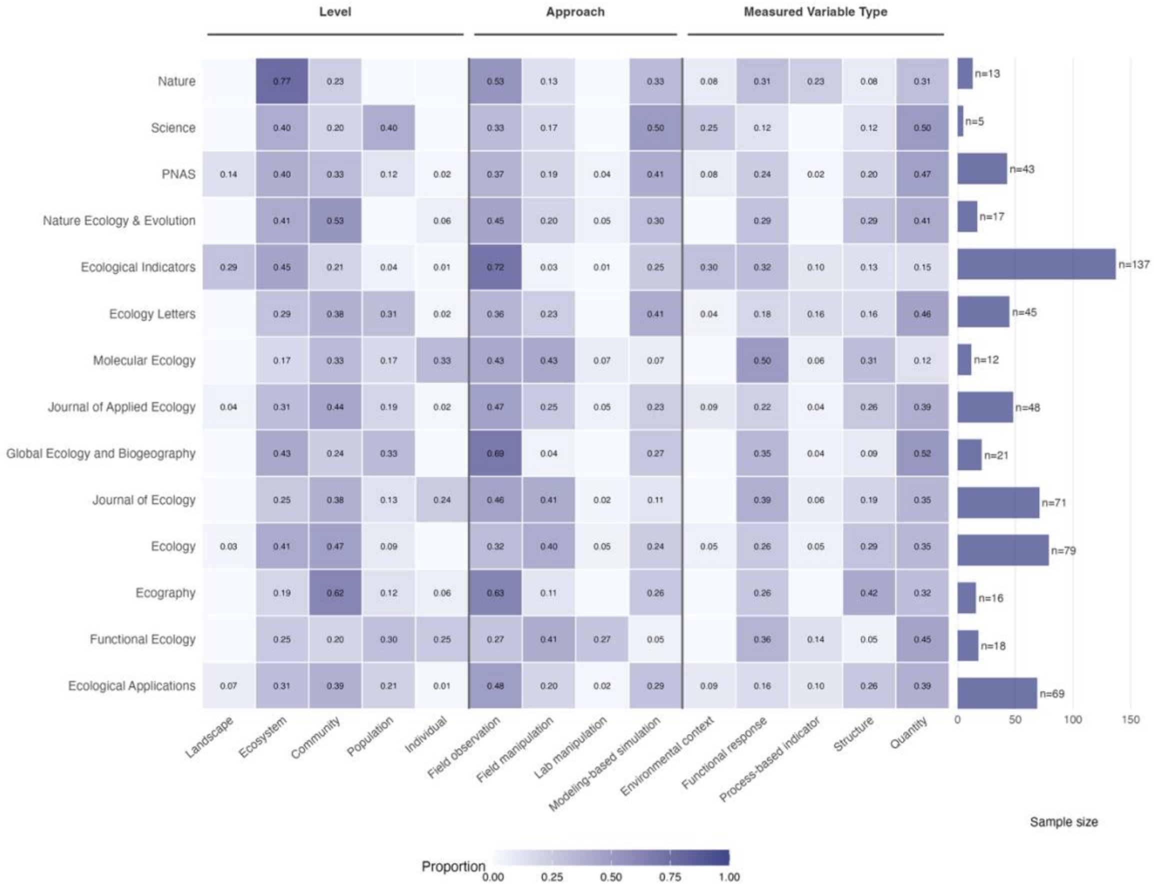


**Supplementary Fig. 11. The distribution of research scope varies substantially across journals.** Heatmap showing the proportion of studies within each journal across ecological level, approach, and measured variable categories. Colour intensity indicates the proportion of studies in each category with a journal, and bar lengths indicate sample size per journal. Journal names are standardised for clarity; in particular, *PNAS* denotes *Proceedings of the National Academy of Sciences of the United States of America*. Although 15 journals were initially selected for this study, Molecular Ecology Resources did not contribute any studies that met the inclusion criteria. As a result, the analyses and this figure include data from 14 journals.

#### S5 Supplementary Information Reference

9. Chen, M., Xu, X., Tan, Y. & Lin, Y. Assessing ecological vulnerability and resilience-sensitivity under rapid urbanization in China's Jiangsu province. *Ecological Indicators* **167**, 112607 (2024).
10. Shi, X. *et al.* Drought risk assessment considering ecosystem resilience: A case study in the Huang-Huai-Hai Plain, China. *Ecological Indicators* **156**, 111102 (2023).
11. Brunswik, E. Organismic achievement and environmental probability. *Psychological review* **50**, 255 (1943).
12. Bender, E. A., Case, T. J. & Gilpin, M. E. Perturbation experiments in community ecology: theory and practice. *Ecology* **65**, 1–13 (1984).
13. IUCN. IUCN Red List categories and criteria: version 3.1. *Prepared by the IUCN Species Survival Commission* (2001).
14. Capdevila, P., Noviello, N., McRae, L., Freeman, R. & Clements, C. F. Global patterns of resilience decline in vertebrate populations. *Ecology Letters* **25**, 240–251 (2022).
15. Hillebrand, H. & Kunze, C. Meta-analysis on pulse disturbances reveals differences in functional and compositional recovery across ecosystems. *Ecology Letters* **23**, 575–585 (2020).
16. Barnett, S. E. & Shade, A. Arrive and wait: Inactive bacterial taxa contribute to perceived soil microbiome resilience after a multidecadal press disturbance. *Ecology Letters* **27**, e14393 (2024).
17. Bonfim, M., López, D. P., Repetto, M. F. & Freestone, A. L. Speed and degree of functional and compositional recovery varies with latitude and community age. *Ecology* **105**, e4259 (2024).
18. Zhang, Y. *et al.* Warming and disturbances affect Arctic-boreal vegetation resilience across northwestern North America. *Nat Ecol Evol* **8**, 2265–2276 (2024).

29. Lever, J. J., Van Nes, E. H., Scheffer, M. & Bascompte, J. Five fundamental ways in which complex food webs may spiral out of control. *Ecology Letters* **26**, 1765–1779 (2023).
30. Blake, C., Barber, J. N., Connallon, T. & McDonald, M. J. Evolutionary shift of a tipping point can precipitate, or forestall, collapse in a microbial community. *Nat Ecol Evol* **8**, 2325–2335 (2024).
31. Power, S. C., Davies, G. M., Wainwright, C. E., Marsh, M. & Bakker, J. D. Restoration temporarily supports the resilience of sagebrush-steppe ecosystems subjected to repeated fires. *Journal of Applied Ecology* **60**, 1607–1621 (2023).
32. Strobl, C., Boulesteix, A.-L., Zeileis, A. & Hothorn, T. Bias in random forest variable importance measures: Illustrations, sources and a solution. *BMC bioinformatics* **8**, 25 (2007).
33. Strobl, C., Boulesteix, A.-L., Kneib, T., Augustin, T. & Zeileis, A. Conditional variable importance for random forests. *BMC bioinformatics* **9**, 307 (2008).
34. Hastie, T., Tibshirani, R. & Friedman, J. The elements of statistical learning. (2009).
35. Liaw, A. & Wiener, M. Classification and regression by randomForest. *R news* **2**, 18–22 (2002).
